# Classical enhancers couple *cis*-regulatory logic with transcriptional condensates and 3D genome architecture

**DOI:** 10.64898/2026.01.23.701252

**Authors:** Ville Tiusanen, Divyesh Patel, Jihan Xia, Chi Xu, Subhamoy Datta, Ji Yun, Samuele Cancellieri, Liangru Fei, Mehmet Yilmaz, Esa Pitkänen, Stefan Prekovic, Päivi Pihlajamaa, Biswajyoti Sahu

## Abstract

Deciphering the regulatory logic of enhancers remains a central question in understanding cell- and tissue-specific gene expression in multicellular organisms. This is particularly pertinent at multipartite enhancer clusters such as super-enhancers, where multiple enhancers contribute to the expression of a single gene. Gene expression has been studied largely through sequence-dependent recruitment of transcription factors (TF) and co-activators, whereas 3D chromatin structure has been attributed to architectural proteins such as cohesion and CTCF^1–3^. However, the contribution of DNA sequence encoded in enhancers to shaping higher-order genome organization remains poorly understood. Here we show that classical enhancers, embedded within multipartite super-enhancer structures, act as determinant regulatory elements that initiate the gene regulatory cascade by linking DNA sequence recognition to 3D chromatin architecture. Classical enhancers are more evolutionarily conserved and display stronger regulatory activity than facilitator elements, which lack intrinsic enhancer activity but potentiate classical enhancer function. We show that classical enhancers are selectively bound by specific TFs with strong intrinsically disordered regions, such as NFE2L2 in liver cancer cells, capable of driving transcriptional condensate formation through phase separation. NFE2L2 depletion reduced enhancer activity and induced widespread chromatin reorganization, characterized by increased cohesin and CTCF occupancy at super-enhancer boundaries and beyond. This “cohesin clogging” impaired DNA loop extrusion, led to formation of smaller topologically associated domains, and weakened enhancer–promoter contacts. These findings highlight that sequence-specific TFs have multifaceted roles beyond transcriptional control, establishing a direct mechanistic link between enhancer sequence, TF binding, condensate formation, and 3D genome organization, with the regulatory logic being encoded in the DNA sequence itself.

## Main

Multicellularity in higher order organisms relies on precise control of cell type-specific gene expression, orchestrated by TFs through *cis*-regulatory elements such as promoters and enhancers^3–5^. Yet the regulatory logic operating at complex, multi-element enhancer clusters – particularly how gene activation is initiated at the level of DNA sequence – remains unresolved.

Gene expression control through typical solitary enhancers is relatively simple, involving TF binding to enhancers that communicate with promoters to activate transcription. In contrast, super-enhancers (SE) are large, multipartite regulatory clusters^6–9^ that control genes central to cellular identity and oncogenic signaling, and their regulatory logic is far more complex. The key question remains: how is the gene expression cascade initiated within these multipartite structures? Does activation emerge from stochastic recruitment of multiple TFs and co-activators, or from an ordered hierarchy in which a specific enhancer and defined TFs trigger the regulatory cascade?

Using functional enhancer activity measurements by self-transcribing active regulatory region sequencing (STARR-seq)^10^, we previously showed that SEs have a multipartite structure comprised of classical enhancers with strong activity in massively parallel reporter assays (MPRA) and chromatin-dependent enhancers that lack intrinsic enhancer activity^11^. Chromatin-dependent enhancers are similar to facilitator elements shown to potentiate classical enhancer function at the α-globin locus^6^. Yet, despite over a decade of SE research^7–9,12–15^, the regulatory logic and functional relevance of their constituent elements remain poorly understood. In particular, it is still elusive whether classical enhancers are defined by unique sequence features or TF binding properties, and whether one distinct enhancer within SEs acts as the primary master regulator to initiate gene expression.

Two complementary mechanisms have emerged to explain enhancer–promoter communication in mammalian genomes: (i) transcriptional condensates formed by phase separation through intrinsically disordered regions (IDR) in co-regulators and RNA polymerase II^16–22^, and (ii) DNA loop extrusion, in which cohesin dynamically folds the genome into 3D structures that facilitate regulatory contacts^23–25^. Some TFs can also drive transcriptional condensate formation via phase separation^21,26–29^, and alterations in TF IDRs through mutations or oncogenic fusions can affect condensate formation, contributing to transcriptional dysregulation in rare genetic disorders^30^ and cancer^31,32^. Condensate formation is especially prominent at SEs^21,28,33^ and has recently been reported to bring the SE in close contact with the gene promoter^34^. However, these models do not explain how sequence-encoded TF binding initiates enhancer hierarchy within SE clusters or how such events affect transcriptional condensate formation and 3D genome architecture. Thus, whether specific TFs selectively bind classical enhancers – rather than other SE elements – to initiate the regulatory cascade remains unknown.

The cohesin complex organizes interphase chromosomes into loops through active extrusion driven by structural maintenance of chromosomes (SMC) proteins^23–25,35,36^. Convergently oriented CTCF sites act as barriers to extrusion^37,38^, creating topologically associating domains (TAD)^39–41^ with frequent intra-domain and limited inter-domain interactions^41,42^. Cohesin loading and unloading controlled by NIPBL and WAPL, respectively^37,43–46^, together with dynamic cohesin turnover, are critical for lineage-specific gene expression^46^. Yet perturbation of CTCF or cohesin often has surprisingly little direct effect on gene expression^47,48^. These observations highlight a major unresolved question: how sequence-specific TF binding at enhancers contributes to loop extrusion dynamics to establish regulatory specificity within SEs. To address this, we combined systematic TF and enhancer perturbations with 3D genome analyses to dissect how the regulatory cascade is initiated at SEs. We show that classical enhancers act as the primary determinant elements within SE clusters, encoding their regulatory logic. This logic is executed through binding of specific TFs with strong IDRs, such as NFE2L2 in liver cancer cells, which initiate the cascade of gene regulatory events, revealing how DNA-encoded regulatory information drives condensate formation and 3D genome organization to activate gene expression.

### Classical enhancers within SEs show selective TF binding

To study the structure and function of classical enhancers within SEs, we mapped 998 SEs in HepG2 liver adenocarcinoma cells using H3K27ac ChIP-seq, with a median length of 16 kb (**Fig. 1a, and Supplementary Table 1**). SEs were enriched for active enhancer mark (H3K27ac), chromatin accessibility (ATAC-seq), the Mediator complex subunit MED1 and enhancer activity measured by STARR-seq (**Fig. 1b**). Based on the STARR-seq signal, we classified the individual enhancers within SEs into classical enhancers (STARR-seq-positive) and chromatin-dependent enhancers^11^ (STARR-seq-negative, henceforth referred to as facilitators^6^) (**Supplementary Table 2**). Representative genome browser snapshot of the *MYC* SE shows strong STARR-seq signal exclusively at the classical enhancer, whereas the adjacent facilitators, by definition, lacked STARR-seq activity (**Fig. 1c**). Notably, both the classical enhancer and the facilitators showed enrichment for ATAC-seq, H3K27ac and MED1 ChIP-seq signals and were distinguished only by the genomic STARR-seq activity, indicating that similar chromatin features do not directly correlate with causal regulatory activity (**Fig. 1c**).

**Fig. 1:**
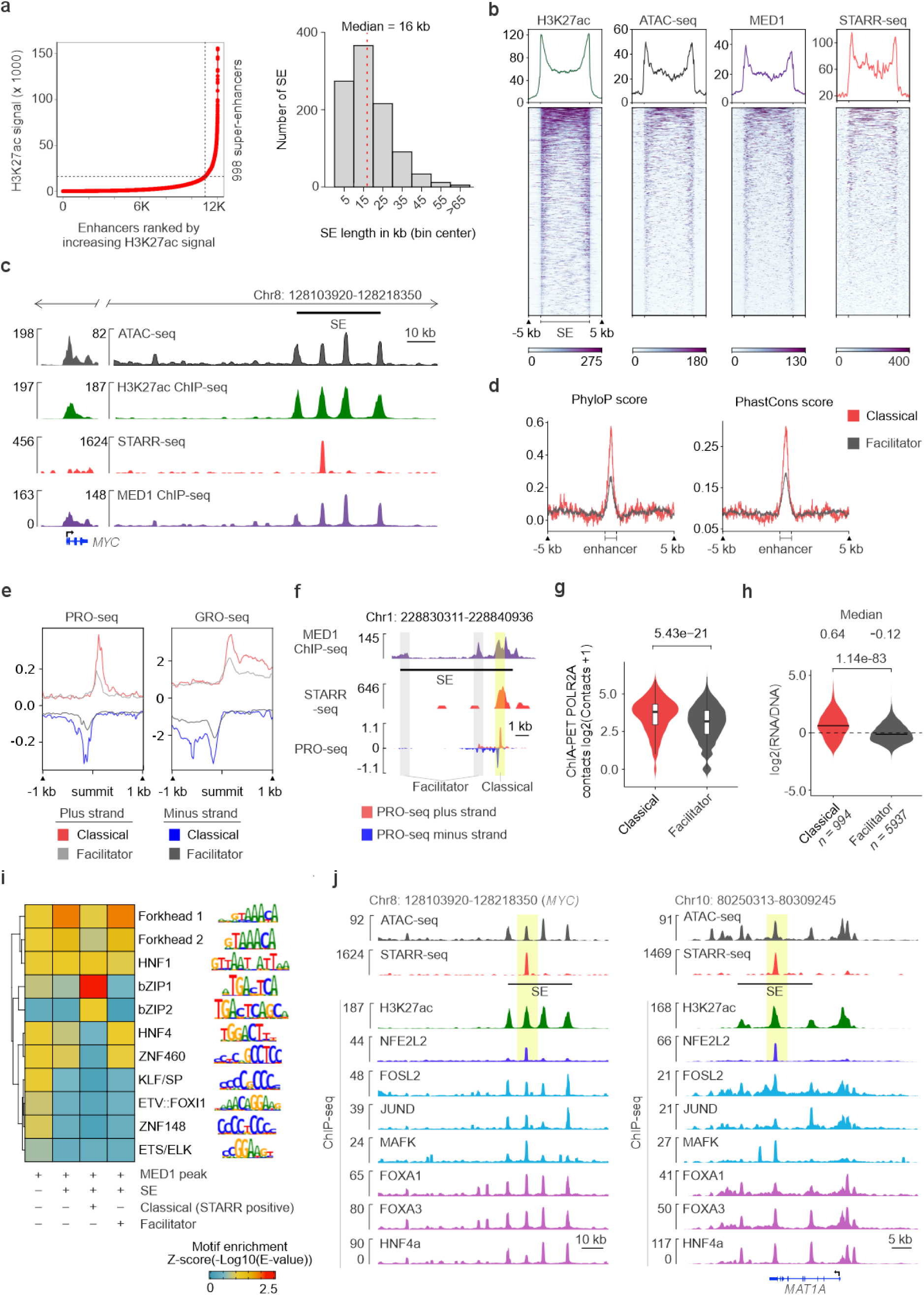
Classical enhancers within SEs are evolutionarily conserved and bound by specific TFs. **a,** Identification of SEs in HepG2 cells based on H3K27ac ChIP-seq signal using ROSE algorithm using stitching distance of 12.5 kb, with ranked candidate enhancers plotted against their H3K27ac ChIP-seq signal (left panel). Histogram showing size distribution of the 998 SEs in the HepG2 cells (right panel). **b,** Heatmap showing RPKM normalized signal for H3K27ac ChIP-seq, ATAC-seq, MED1 ChIP-seq and STARR-seq at HepG2 SEs, plotted for each entire SE region including ±5 kb flanking regions. **c,** Genome browser snapshot from the *MYC* locus showing the downstream SE consisting of one classical enhancer (marked by STARR-seq signal) and three facilitators. Both classical enhancer and facilitators are enriched for ATAC-seq, and ChIP-seq signal for MED1 and H3K27ac. **d,** Comparison of evolutionary conservation score between classical enhancers (n = 496) and facilitators (n = 3476) in HepG2 cells, shown as metaplots comparing PhyloP100 (left) and PhastCons100 (right) scores. **e,** Metaplot showing strand-specific PRO-seq and GRO-seq signal for classical enhancers (n = 496) and facilitators (n = 3476). **f,** Genome browser snapshot showing PRO-seq signal at SEs. Classical enhancers (highlighted in yellow) exhibit stronger nascent transcriptional activity compared to facilitators (highlighted in grey). Each panel shows MED1 ChIP-seq (purple), STARR-seq (red), and PRO-seq (blue) signals. **g,** Comparison of number of interactions between classical enhancers (n = 496) and facilitators (n = 3476) measured by POLR2A ChIA-PET (unpaired Wilcoxon test). Boxes represent median and interquartile range. **h,** Classical enhancers show consistently higher enhancer activity than facilitators under both episomal and chromatinized conditions. Violin plots showing the log2(RNA/DNA) ratios from the HepG2 lenti-MPRA data^51^ overlapping with classical enhancers and facilitators determined using STARR-seq data (unpaired Wilcoxon test). Solid line represents median value for each group. **i,** TF motif enrichment analysis using Analysis of Motif Enrichment (AME)^53^ for four groups of enhancers: i) enhancers outside SEs, ii) enhancers inside SEs, iii) classical enhancers within SEs, iv) facilitators within SEs. After performing motif enrichment analysis for individual motifs, similar motifs were combined into motif clusters according to Viestra et al. 2020^54^. The representative TF families and motifs are shown on the right. **j,** Classical enhancers within HepG2 SEs are selectively bound by NFE2L2. Genome browser snapshots showing NFE2L2 binding at classical enhancers within SEs on chromosome 8 (left) and *MAT1A* locus (right) in HepG2 cells. Each panel shows signal for ATAC-seq (black), STARR-seq (red), and ChIP-seq signal for H3K27ac (green), NFE2L2 (dark blue), bZIP family TFs (light blue), and other TFs (purple).

Classical enhancers within SEs showed greater evolutionary conservation than facilitators (**Fig. 1d**), consistent with stronger selective pressure on their functional importance. They were also more enriched for H3K27ac (**Extended Data Fig. 1a**) and exhibited higher levels of divergent enhancer RNA transcription, measured by using precision run-on sequencing (PRO-seq) and global run-on sequencing (GRO-seq)^49^ (**Fig. 1e-f and Extended Data Fig 1b**). Chromatin immunoprecipitation–paired-end tag (ChIA-PET) analysis using POLR2A in HepG2 cells^50^ further revealed that classical enhancers form significantly more genomic interactions than facilitators (**Fig. 1g**), underscoring their regulatory importance. Moreover, recently reported lenti-MPRA data in HepG2 cells^51^ allowed comparison of enhancer activity between our episomal STARR-seq assay and an orthogonal assay in native chromatin context. Importantly, lenti-MPRA fragments overlapping with classical enhancers showed strong activity, whereas those overlapping with facilitators had a negative median log2(RNA/DNA) ratio (**Fig. 1h and Extended Data Fig. 1c**). These results confirm that classical enhancers can be robustly identified using STARR-seq, demonstrating that STARR-seq – an episomal (‘out-of-genome’) readout – faithfully captures activity of classical enhancers, despite lacking the fully chromatinized environment.

We asked whether the functional differences between classical enhancers and facilitators could be explained by their underlying sequence composition. To test this, we performed TF motif enrichment analysis that can discriminate between these two types. Enhancers overlapping with SEs encompassing both classical enhancers and facilitators were enriched for motifs from the forkhead box (FOX), hepatocyte nuclear factor (HNF) and basic leucine zipper (bZIP) TF families compared to non-SE enhancers. Interestingly, classical enhancers within SEs showed a strong enrichment for motifs of the bZIP family TFs, particularly NFE2L2 (TGAnTCAGCA) (**Fig. 1i, and Supplementary Table 3**). Consistently, TF affinity prediction using a biophysical model (TRAP)^52^ showed that bZIP family TFs have a stronger binding preference for classical enhancers than for facilitators (**Extended Data Fig. 1d and Supplementary Table 4**). Leveraging the ENCODE ChIP-seq datasets in HepG2 cells^50^, we confirmed that NFE2L2 selectively binds to classical enhancers, whereas facilitators were bound by other TFs identified from motif analysis, such as FOXAs, HNF4a, as well as other, non-selective, bZIP-family TFs (**Fig. 1j and Extended Data Fig. 1e**). These results suggest that NFE2L2 is a key regulator of classical enhancers in HepG2 cells.

To assess whether these observations extend beyond HepG2 cells, we analyzed classical enhancers and facilitators in GP5d colon adenocarcinoma cell line. Using H3K27ac ChIP-seq, we identified 1319 SEs with a median size of 24 kb (**Extended Data Fig. 2a and Supplementary Table 1**). These SEs were enriched for STARR-seq, ATAC-seq and MED1 ChIP-seq signals (**Extended Data Fig. 2b**), and classical enhancers defined by STARR-seq (**Extended Data Fig. 2c and Supplementary Table 2**) showed stronger evolutionary conservation than facilitators (**Extended Data Fig. 2d**). Motif enrichment analysis revealed distinct TF preferences compared to HepG2 cells. SE-overlapping enhancers in GP5d were enriched for CDX, FOX, and SNAI/TCF family motifs (**Extended Data Fig. 2e and Supplementary Table 3**). Notably, classical enhancers were specifically enriched for TCF7 motif, whereas facilitators were enriched for FOXA motifs, resembling the pattern observed in facilitators in HepG2 cells (**Extended Data Fig. 2e**). TF affinity prediction further indicated that TCF7L2 preferentially binds classical enhancers (**Extended Data Fig. 2f, and Supplementary Table 4**), and ChIP-seq data confirmed selective occupancy of TCF7L2 at these sites (**Extended Data Fig. 2g**). Collectively, these results demonstrate that classical enhancers are functionally distinct from facilitators, and that they are preferentially bound by distinct TFs in a cell type-selective manner.

### NFE2L2 IDR drives biomolecular condensate formation

Given that NFE2L2 selectively binds classical enhancers, the primary enhancers within SEs, we hypothesized that it could mediate transcriptional condensate formation at SEs, bringing together classical enhancers and the facilitators that lack intrinsic enhancer activity. Structural prediction using AlphaFold 3^55^ revealed that NFE2L2 contains a long region with low confidence, which correlates to predicted IDR^56^ (**Fig. 2a**). Moreover, sequence-based IDR analysis identified a stretch of polar and charged amino acids between residues 333–456, immediately upstream of its DNA-binding domain (**Fig. 2b**). In contrast, several other bZIP factors such as MAFF, MAFK, and ATF3 lacked defined IDRs (**Fig. 2b and Extended Data Fig**. **3a**). Similarly, TCF7L2, the classical enhancer-specific TF in GP5d cells, also contained a strong IDR, suggesting a similar preference for classical enhancers in SE-mediated gene regulation (**Extended Data Fig. 3a**).

**Fig. 2:**
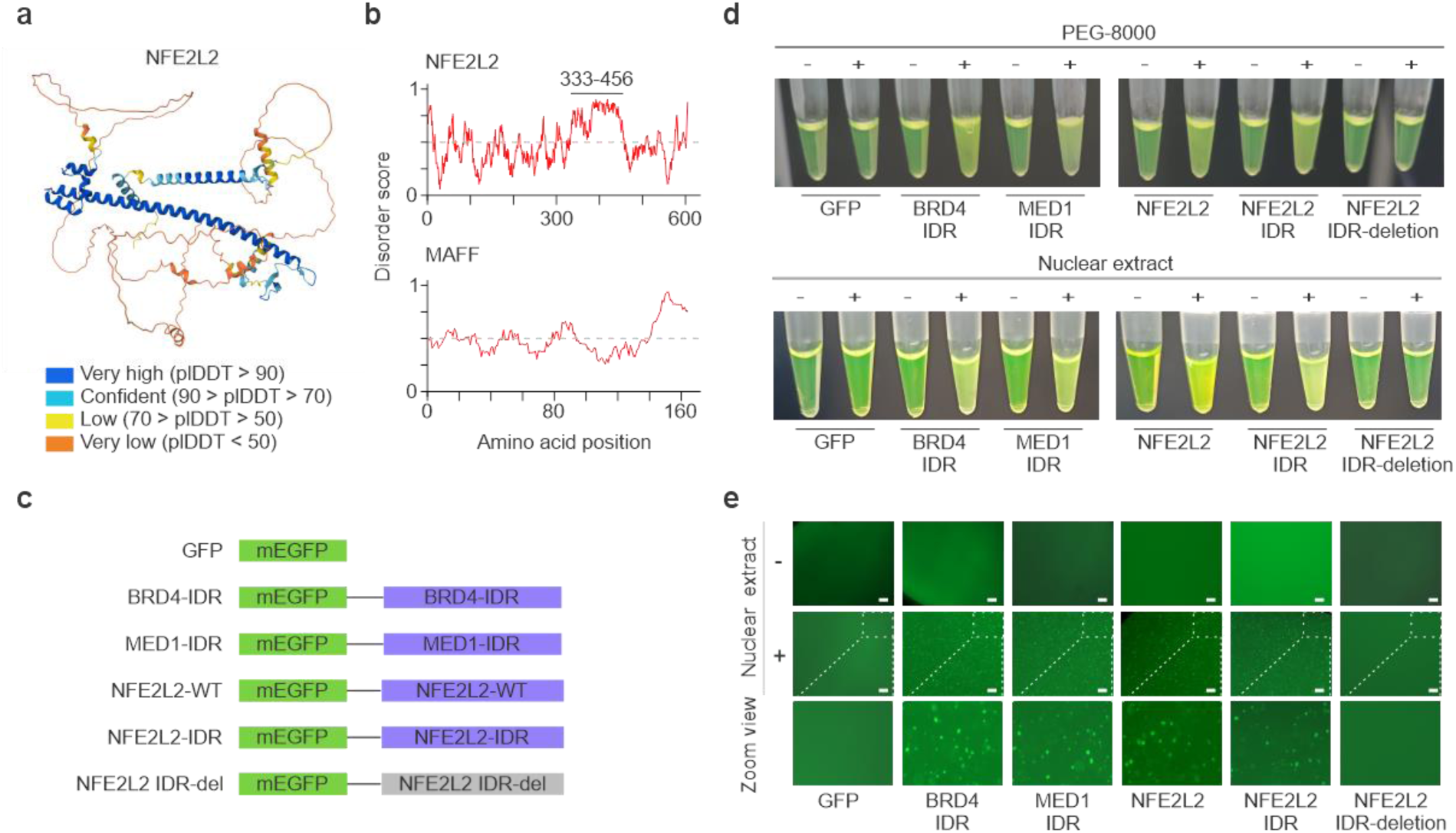
NFE2L2 IDR drives phase separation in vitro. **a,** AlphaFold 3^55^ prediction of the NFE2L2 protein structure (Uniprot: Q16236). Predicted local distance difference test (pLDDT) values (indicated with colors) correspond to accuracy of the prediction, regions with low confidence (orange, yellow) correlating to predicted IDR. **b,** Sequence-based IDR prediction for NFE2L2 (top) and MAFF (bottom) proteins by using IUPred2A^58^. Y-axis represents the disorder prediction score plotted for each amino acid position. Scores over 0.5 (gray dashed line) are considered disordered. Amino acid residues forming IDR in NFE2L2 are highlighted with line, whereas MAFF does not contain predicted IDRs. **c,** Schematic illustrations of the constructs used for phase-separation assays. Constructs with intact IDR domains are marked with blue and IDR-deficient NFE2L2 construct is highlighted in grey. **d,** *In vitro* turbidity assay for phase separation. From left to right: tubes containing GFP (negative control), BRD4-IDR (positive control), MED1-IDR (positive control), NFE2L2 WT, NFE2L2 IDR, and IDR-deficient NFE2L2 constructs in the presence (+) or absence (−) of PEG-8000 (top) or nuclear extract (bottom). **e,** Representative microscopic images of droplet formation in the presence of nuclear extract. Bottom row shows zoomed-in view of the microscope images. All scale bar is 50 µM.

To test whether NFE2L2 can induce phase separation through its IDR, we purified mGFP-tagged fusion proteins including BRD4-IDR and MED1-IDR as positive controls with established phase separation activity^21,29^, along with full-length NFE2L2, NFE2L2-IDR, and an IDR-deficient NFE2L2 variant (**Fig. 2c**). In the presence of 10% PEG8000 or nuclear extract, turbidity assay showed that BRD4-IDR, MED1-IDR, NFE2L2, and NFE2L2-IDR formed phase-separated condensates, whereas the IDR-deficient NFE2L2 did not (**Fig. 2d**). Consistently, recombinant NFE2L2 and NFE2L2-IDR also formed droplets in presence of nuclear extract (**Fig. 2e**). Collectively, these results indicate that the IDR of NFE2L2 is sufficient to drive phase separation, consistent with its annotation as a phase separation–associated TF in the CD-CODE database^57^. This suggests a mechanism for how NFE2L2 binding at classical enhancers can induce transcriptional condensates that coalesce both classical enhancers and facilitators within SE clusters.

### NFE2L2 loss reduces H3K27ac levels across SEs and increases CTCF and SMC1 binding at the SE flanks

To investigate the role of NFE2L2 in the regulation of SEs, we depleted NFE2L2 in HepG2 cells using CRISPR-Cas9 (NFE2L2 knockdown, KD) (see Methods for details; **Extended Data Fig. 3b-c**). NFE2L2 ChIP-seq confirmed efficient knockdown with a 95% reduction in peak number and markedly decreased signal intensity (**Fig. 3a-c**). Transcriptional profiling revealed widespread changes, with 1,318 differentially expressed genes upon NFE2L2 depletion (**Extended Data Fig. 3d**).

**Fig. 3:**
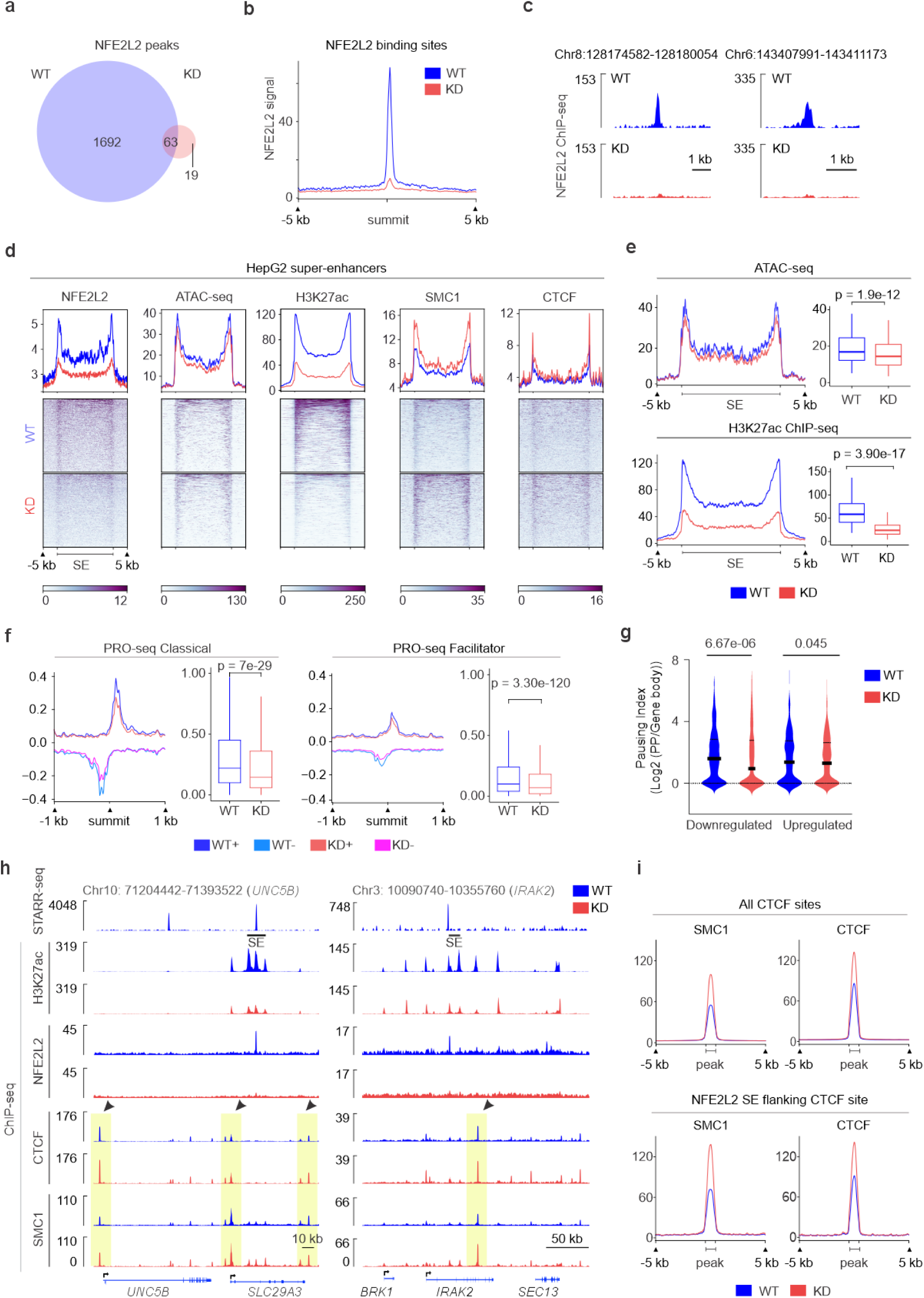
NFE2L2 knockdown reduces H3K27ac levels across SEs and increases CTCF and SMC1 binding at the SE flanks. **a,** Venn diagram showing overlap of NFE2L2 ChIP-seq peaks in HepG2 WT (blue) and NFE2L2-depleted cells (KD; red). Reproducible peaks present in two replicates were used for overlap analysis. **b,** Metaplot comparing NFE2L2 ChIP-seq signal from HepG2 WT and HepG2 NFE2L2-depleted cells. RPKM-normalized ChIP-seq signal was plotted in 5 kb flanks from the center of the NFE2L2 ChIP-seq peaks for HepG2 WT cells (n = 1755). **c,** Genome browser snapshots for NFE2L2 ChIP-seq signal. Each panel shows RPKM-normalized NFE2L2 ChIP-seq signal for HepG2 WT and NFE2L2-depleted cells. **d,** Heatmap showing signal for NFE2L2 ChIP-seq, ATAC-seq, ChIP-seq for H3K27ac, SMC1 and CTCF at all HepG2 SEs (n=998) from HepG2 WT (blue) and HepG2 NFE2L2-depleted cells (red), RPKM-normalized signals plotted for each entire SE region including ±5 kb flanking regions. **e,** Metaplots comparing signal for ATAC-seq (upper panel) and H3K27ac ChIP-seq (bottom panel) at SEs harboring a NFE2L2-bound classical enhancer (n=87). Boxplots showing mean RPKM-normalized signals for ATAC-seq and H3K27ac ChIP-seq in WT and NFE2L2-depleted HepG2 cells (paired Wilcoxon test). Boxplots display the median and interquartile range. **f,** Metaplots comparing strand-specific PRO-seq signal at classical enhancers (left) and facilitators (right) in HepG2 WT (dark & light blue) and NFE2L2-depleted cells (KD; red and pink); boxplots showing quantification of absolute value of mean PRO-seq signals (paired Wilcox test). Boxplots display the median and interquartile range. **g,** Violin plots showing RNA Pol II pausing index analyzed using PRO-seq (see Supplementary Methods for details) for differentially expressed genes between HepG2 WT and NFE2L2-depleted cells (downregulated, n= 801; upregulated, n= 367 from RNA-seq data with cutoff |Log2FC|>1.5 and FDR 0.05; paired t-test, two-sided). Median indicated with continuous line, the first and third quartiles with dashed line. **h,** Genome browser snapshot showing SEs at *UNC5B* (left) and *IRAK2* locus (right). Each panel shows STARR-seq signal in WT (blue), and ChIP-seq signal for NFE2L2, H3K27ac, CTCF and SMC1 in both WT (blue) and NFE2L2-depleted cells (red). Arrowhead indicate CTCF binding sites, showing gain of ChIP-seq signal for CTCF and SMC1 in NFE2L2-depleted cells. **i,** Metaplots comparing ChIP-seq signal for SMC1 and CTCF at all CTCF-binding sites (upper panel) and CTCF-binding sites flanking the SEs harboring NFE2L2-bound classical enhancers (lower panel). RPKM-normalized ChIP-seq signal was plotted for 5 kb regions around the CTCF ChIP-seq peak.

Given the strong enrichment of NFE2L2 at classical enhancers, we expected its depletion to induce strong changes in chromatin accessibility and active enhancer status with reduced coactivator binding. Instead, analysis of all SEs (n=998) showed no appreciable reduction in chromatin accessibility or in BRD4 occupancy. Intriguingly, H3K27ac levels were strongly reduced, concomitant with marked reduction in NFE2L2 binding but with no change in p300 binding (**Fig. 3d and Extended Data Fig. 4a**). However, at SEs harboring NFE2L2-bound classical enhancers (n=87), we observed a modest but significant reduction in chromatin accessibility along with pronounced decrease in H3K27ac (**Fig. 3e**). NFE2L2 knockdown also resulted in stronger reduction in nascent enhancer RNA transcription at classical enhancers compared to facilitators (**Fig. 3f and Extended Data Fig. 4b**). Consistently, PRO-seq revealed reduced RNA Pol II recruitment at downregulated genes in NFE2L2-depleted cells (**Fig. 3g**).

The observation that NFE2L2 depletion resulted only in reduction of H3K27ac levels both at SEs and genome-wide but with little to no effect on p300 and BRD4 occupancy or chromatin accessibility, led us to look for changes in chromatin structure. Notably, NFE2L2 depletion led to increased binding of the architectural proteins CTCF and SMC1, particularly at genomic regions immediately flanking the SEs (**Fig. 3d**). To avoid clone-specific bias, we confirmed similar changes in chromatin accessibility, H3K27ac levels and CTCF occupancy in another NFE2L2 knockdown clone (KD2; **Extended Data Fig. 3b-c and Extended Data Fig. 4c**). The increased binding of CTCF and SMC1 extended beyond the SEs at broader genomic intervals in an almost symmetrical fashion, as illustrated for the *UNC5B* and *IRAK2* loci (**Fig. 3h**). Increased CTCF and SMC1 occupancy was observed at all genome-wide CTCF sites and particularly at CTCF sites flanking the NFE2L2-bound SEs (**Fig. 3i**). One possible explanation for this increased loading could be deregulation of cohesin loading and unloading proteins, NIPBL and WAPL, respectively. However, the binding of these proteins was largely unaffected after NFE2L2 depletion (**Extended Data Fig. 4d**). Collectively, these results indicate that NFE2L2 depletion reduces classical enhancer activity and alters the binding pattern of chromatin architectural proteins, with increased CTCF and SMC1 binding at SE flanks and beyond.

### NFE2L2 depletion reshapes chromatin architecture and enhancer-promoter interactions

The increased CTCF and SMC1 binding upon NFE2L2 depletion at SE boundaries and at regions flanking the SEs in a symmetrical pattern suggests reorganization of 3D chromatin structure, possibly due to impaired DNA loop extrusion. Polymer modeling simulations using MoDLE^59^ (see Methods for details) revealed a shift in TAD size distribution toward smaller domains and a more negative TAD insulation score, indicative of enhanced insulation (**Fig. 4a**). The number of predicted TADs based on CTCF ChIP-seq data increased from 8,880 in WT to 9,327 in NFE2L2-depleted cells, suggesting that loss of classical enhancer-bound NFE2L2 disrupts productive enhancer activity at SEs, leading to weaker or stalled loop extrusion.

**Fig. 4:**
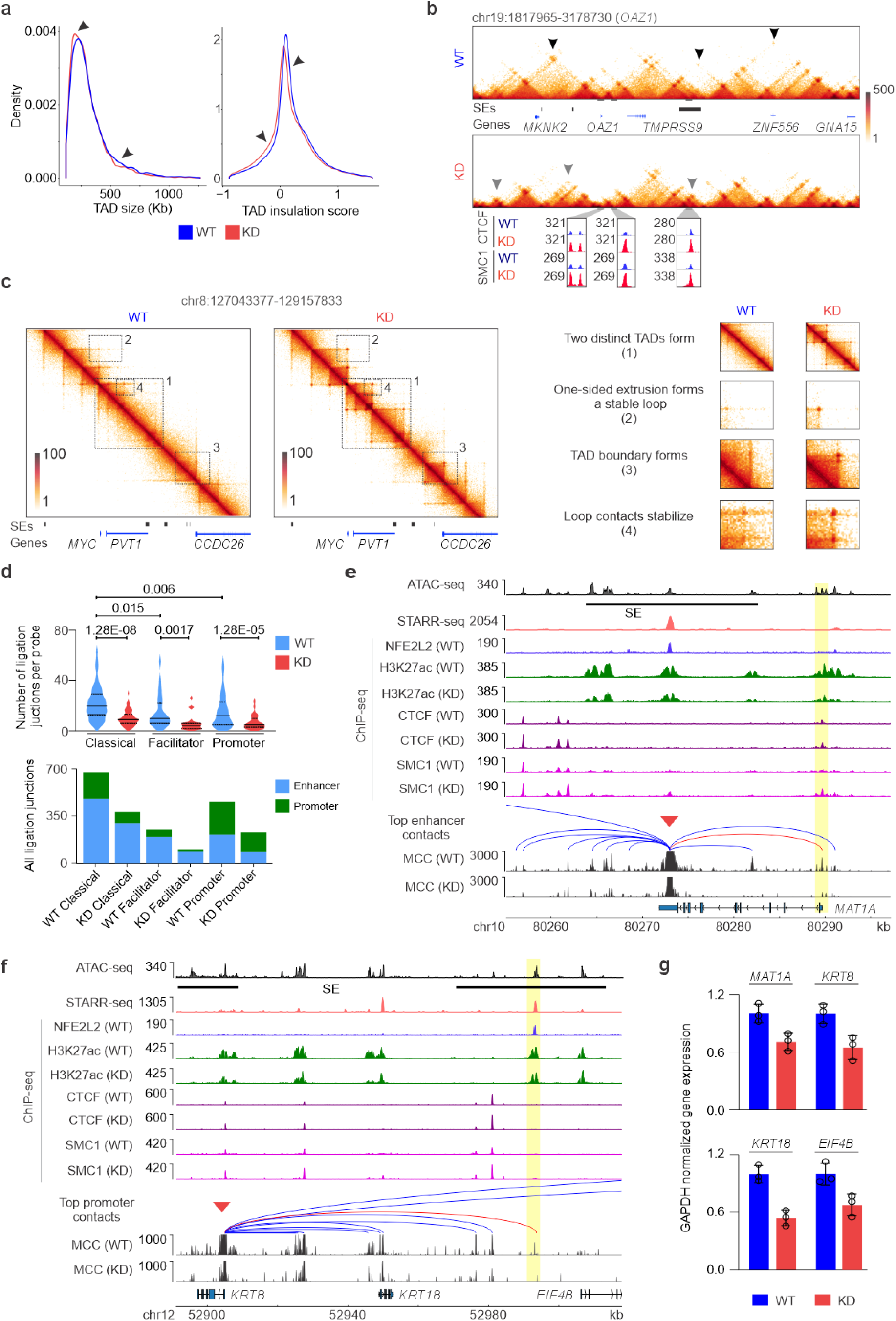
NFE2L2 depletion reshapes chromatin architecture and enhancer-promoter interactions. **a,** Density plots showing the distribution of TAD lengths (left) and insulation scores at TAD boundaries (right) by using simulated genome wide loop extrusion contacts for HepG2 WT and NFE2L2-depleted cells. Polymer simulation was performed at resolution of 20 kb using MoDLE^59^ (see Methods for details). **b,** Comparison of loop extrusion contacts frequency at *OAZ1* locus between HepG2 WT and NFE2L2-depleted cells. Top panel shows the predicted loop extrusion contacts using CTCF ChIP-seq at *OAZ1* gene locus. The bottom panel shows ChIP-seq signal for CTCF and SMC1 in HepG2 WT (blue) and NFE2L2-depleted cells (red) at the marked regions. **c,** Left panel: simulated loop extrusion contacts using CTCF ChIP-seq for *MYC* locus in HepG2 WT (left) and NFE2L2-depleted cells (right). A larger TAD comprising *MYC* gene and nearby SEs sub-divided in multiple smaller sub-TADs upon NFE2L2-depletion. Right panel: simulated genome-wide loop extrusion data reveal distinct chromatin alterations by NFE2L2-depletion, including: (i) fragmentation of a large TAD into smaller sub-TADs (top), (ii) stabilization of a one-sided loop extrusion into a stable loop (upper middle), (iii) emergence of a new TAD boundary (lower middle), and (iv) stabilization of loop contacts (bottom). **d,** Top: violin plot showing the number of ligation junctions per probe for classical enhancers, facilitators and promoter MCC viewpoints in WT and NFE2L2-depleted cells (unpaired t-test, two-sided). Solid and dashed lines represent median and interquartile range, respectively. Bottom: genomic annotation for the ligation junction contacts for classical enhancers, facilitators and promoter viewpoints for WT and NFE2L2-depleted cells. **e,** MCC contact profile for the NFE2L2-bound classical enhancer at the *MAT1A* locus (viewpoint at NFE2L2-bound classical enhancer). Each panel shows ATAC-seq, STARR-seq and NFE2L2 ChIP-seq for HepG2 WT cells and ChIP-seq for H3K27ac, CTCF and SMC1 for HepG2 WT and NFE2L2-depleted cells. DNA loop between classical enhancer and *MAT1A* promoter is highlighted in red. **f,** MCC contact profile for the NFE2L2-bound classical enhancer at the *KRT8* locus (viewpoint at NFE2L2-bound classical enhancer). Panel description same as Fig. 4e. DNA loop between classical enhancers and *KRT8* promoter is highlighted in red. **g,** RT-qPCR data showing changes in mRNA expression for selected SE-target genes in WT and NFE2L2 -depleted cells. The GAPDH normalized expression for each gene were compared relative to HepG2 WT cells. The figures show mean ± SD values for three technical replicates.

To illustrate these genome-wide changes at the locus level, we examined representative SE-associated regions. For example, near the *OAZ1* gene, simulations based on both CTCF (**Fig. 4b**) and SMC1 ChIP-seq data (**Extended Data Fig. 5a**) revealed the formation of new TADs in NFE2L2-depleted cells. As CTCF more accurately predicts loop extrusion dynamics compared to SMC1^59^, we used CTCF ChIP-seq signal for these structural predictions. Upon NFE2L2 depletion, larger TADs were frequently disrupted into sub-TADs near SEs associated with NFE2L2-bound classical enhancers (**Extended Data Fig. 5b-c**). We next examined the predicted structural changes at a well characterized genomic region harboring the *MYC* oncogene^60,61^. In WT HepG2 cells, the *MYC* gene and its downstream SEs reside within a large TAD (**Fig. 4c**, left panel). In NFE2L2-depleted cells, this TAD was subdivided into multiple smaller sub-TADs, resulting in increased insulation between the *MYC* promoter and downstream SEs. We identified several distinct chromatin remodeling events at the *MYC* locus: (i) subdivision of a large TAD into multiple sub-TADs, (ii) one-sided loop extrusion forming stable loops, (iii) emergence of new TADs, and (iv) stabilization of chromatin loop contacts (**Fig. 4c**, right panel). These new sub-TAD boundaries coincided with increased CTCF and SMC1 occupancy, suggesting a mechanistic link to altered chromatin interactions. Polymer simulation using either CTCF or SMC1 ChIP-seq data as input predicted similar reorganization at the *MYC* locus (**Extended Data Fig. 5d-e**).

To directly investigate enhancer–promoter interactions at base-pair resolution, we performed micro-capture-C (MCC)^62^ targeting NFE2L2-bound classical enhancers and gene promoters in WT and NFE2L2-depleted HepG2 cells. We compared ligation junctions across distinct categories of MCC bait-regions and found that MCC probes at classical enhancers showed a higher number of interactions than those linked to facilitators or promoters in both WT and NFE2L2-depleted cells (**Fig. 4d, top panel; and Extended Data Fig. 6a**), suggesting their dominant role as primary enhancers in the SE cluster. Similarly, MCC probes at classical enhancers had higher interaction frequency with other enhancer elements including the facilitators, whereas probes at facilitators showed comparatively fewer interactions in both WT and NFE2L2-depleted cells (**Fig. 4d).** Formation of DNA loops between classical enhancers and promoters was further confirmed using independent probes targeting both elements at the same locus (**Extended Data Fig. 6b**). Upon NFE2L2 depletion, classical enhancer-promoter interactions were markedly reduced for genes downregulated in KD cells (**Fig. 4e-f and Extended Data Fig. 6c**), and decreased expression of these genes was validated using RT–qPCR (**Fig. 4g**). In addition to enhancer-promoter contacts, enhancer–enhancer interactions between SE-associated enhancers and nearby regulatory elements were also diminished in NFE2L2-depleted cells (**Extended Data Fig. 6d**), consistent with reduced H3K27ac and PRO-seq signals at SEs (c.f. **Fig. 3e-f**). Together, these results demonstrate that NFE2L2 depletion leads to increased TAD formation and stronger domain insulation, disrupting enhancer–promoter interactions.

### Classical enhancers exhibit stronger cis-regulatory activity than facilitators

To determine the enhancer potential of classical enhancers and facilitators within SEs, we performed CRISPR-Cas9-mediated deletions using guide RNA pairs flanking each individual element (**Extended Data Fig. 7a**). Homozygous deletion clones were generated for classical enhancers and/or facilitators across four SEs, and transcript levels of candidate target genes were quantified by RT-qPCR. At the *KRT8/KRT18* and *MKKS* loci (**Fig. 5a**), deletion of classical enhancers caused markedly stronger reductions in target gene expression than deletion of facilitator elements (**Fig. 5b and Extended Data Fig. 7b**). For example, classical enhancer deletion reduced *KRT8* and *KRT18* expression by 39% and 19%, respectively, compared with 9–11% and 1–4% reductions following facilitator deletions. Similarly, classical enhancer deletion reduced expression of the *FN1* and *ERRFI1* genes by more than 50% and 30%, respectively (**Fig. 5b and Extended Data Fig. 7b**).

**Fig. 5:**
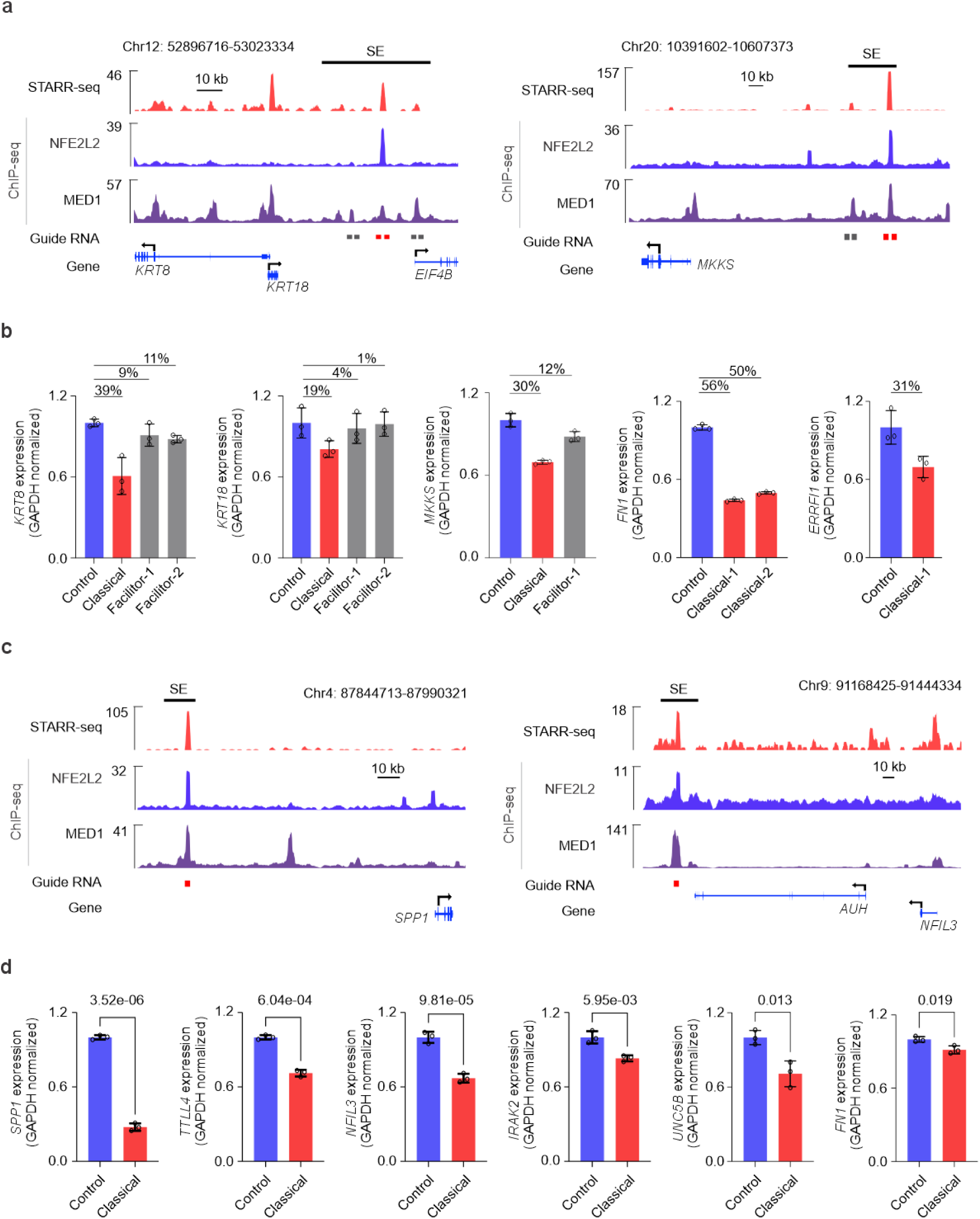
Classical enhancers exhibit stronger cis-regulatory activity than facilitators. **a,** CRISPR-Cas9-mediated deletion of classical enhancers and/or facilitators within SEs in HepG2 cells. Representative genome browser views for *KRT8* and *MKKS* loci showing sgRNAs targeting each site of the classical enhancers, along with STARR-seq and ChIP-seq signals for NFE2L2 and MED1 in HepG2 cells. **b,** RT-qPCR data showing changes in mRNA expression for target genes upon CRISPR-Cas9 mediated deletion of classical enhancers and/or facilitators within SEs in HepG2 cells. GAPDH normalized expression were compared relative to control cells. The figures show mean ± SD values for three technical replicates. **c,** CRISPR interference (CRISPRi) using dCas9-KRAB-MeCP2 targeting the summits of classical enhancers. Representative genome browser views for *SPP1* and *NFIL3* loci showing sgRNAs targeting the summits of the classical enhancers of the SEs. **d,** RT-qPCR data showing changes in mRNA expression for target genes upon dCas9-KRAB-MeCP2 mediated silencing of classical enhancers within SEs in HepG2 cells. GAPDH normalized expression were compared relative to control cells. The figures show mean ± SD values for three technical replicates.

We further validated enhancer function using CRISPR interference (CRISPRi) with dCas9-KRAB-MeCP2 and a single guide RNAs (sgRNA) targeting the summits of classical enhancers within six SEs, including the *SPP1* and *NFIL3* loci (**Fig. 5c**). In all six cases, silencing of the classical enhancer reduced expression of the target gene compared to control cells as measured by RT-qPCR. (**Fig. 5d**). Together, these results demonstrate that classical enhancers on their own have higher potential in activating gene expression compared to facilitators within SEs.

Taken together, we show that genomic STARR-seq enables faithful identification of classical enhancers from the facilitators within the SEs. Classical enhancers act as deterministic enhancer elements within the multi-partite enhancers like SEs, and are bound by specific TFs with strong IDRs that nucleate facilitators into transcriptional condensates to promote enhancer-promoter interaction within proper 3D chromatin structure. Depletion of classical enhancer-specific TF such as NFE2L2 disrupts phase separation, diminishes enhancer-associated features, and reorganizes chromatin architecture, thereby impairing enhancer-promoter interactions and leading to transcriptional dysregulation showing the regulatory logic of DNA sequence linking the full cascade from sequence-specific TF binding to 3D genome organization.

## Discussion

Deciphering the regulatory logic of enhancers remains a central question in understanding cell- and tissue-specific gene expression in multicellular organisms. This locus-specific and long-distance control of gene expression is particularly complex in the case of multiple clustered enhancers as in SEs compared to typical solitary enhancers. Large-scale enhancer activity assays such as STARR-seq^10,11,63–65^ and lenti-MPRA^51,66^ combined with deep learning^67–70^ have begun to illuminate enhancer grammar and enabled rational design of synthetic enhancers^71,72^ with applications in basic biology and therapeutic development^71–74^. However, productive gene expression is a complex process involving sequence-specific TF binding, recruitment of co-regulators, and physical interactions between enhancers and promoters. How this process is initiated and hierarchically coordinated within complex regulatory architectures—such as multipartite enhancer clusters and SEs where multiple elements converge on the same target gene—remains poorly understood and is difficult to envision as a random process of sequestration of multiple TFs and coactivators. Thus, a fundamental unresolved question in enhancer biology is, how regulatory logic encoded within the DNA sequence determines which enhancer acts as the primary driver of gene regulatory cascade when multiple enhancers are involved in complete activation of a given gene.

Here, we identify a previously unrecognized class of TFs, “lexifiers”, that decode the regulatory lexicon embedded within enhancer sequences to initiate gene expression cascade. The classical enhancers that we envision as “primal enhancers” within multipartite enhancer clusters recruit the lexifier TF, which interprets the enhancer grammar and induces the formation of transcriptional condensates through phase separation. Disruption of this lexifier–primal enhancer axis leads to impaired chromatin loop dynamics and accumulation of architectural proteins at SE boundaries – a phenomenon we term “cohesin clogging”. Thus, our results establish a direct mechanistic link between sequence-specific TF binding, phase separation and 3D chromatin organization, where the regulatory logic is encoded in the DNA sequence itself and is deciphered by the lexifier TFs.

Multipartite SEs are characterized by high concentrations of TFs and co-regulators that promote the formation of phase-separated transcriptional condensates^21^, which dynamically engage in enhancer–promoter interactions described by the “three-way kissing” model^34^. Thus, SEs provide an ideal spatial framework to resolve the hierarchical logic of multipartite enhancer assemblies. However, beyond a few well-studied loci such as α-globin^6,12^ and SOX^75^, the contribution of individual elements has remained elusive along with the ordered hierarchy of these elements. Here, we show that classical enhancers act as the primary and fundamental regulatory elements within SEs, recruiting the lexifier TFs to translate the genomic lexicon into higher-order transcriptional organization. Classical enhancers are more evolutionarily conserved than facilitators, thus termed as primal enhancers, and display higher enhancer activity in both episomal and chromatin contexts, as shown by STARR-seq and lenti-MPRA data^51^. Functional perturbations using CRISPR-Cas9 further validated classical enhancers as the critical regulatory nodes within SEs. Facilitators, on the other hand, contribute to the expression of the same genes by potentiating classical enhancer activity, consistent with observations at the α-globin locus^6^. Together, these results support a weak enhancer syntax^76^, in which gene expression output reflects the cumulative contribution of multiple regulatory elements but is hierarchically initiated by one primal enhancer as seen in SEs.

Lexifier TFs that decode the regulatory logic at SEs share two defining properties: selective binding to classical enhancers and the ability to drive phase separation through strong IDRs at physiologically relevant expression levels, with higher abundance typically enhancing condensate formation^21^. Our results indicate that lexifier TFs are cell type-specific, such as NFE2L2 in liver cancer and TCF7L2 in colon cancer cells, but whether other TFs with strong IDRs and activation domains can function in similar roles remains to be investigated. Notably, other bZIP TFs expressed in HepG2 cells, such as JUN and FOS, also contain IDRs but bind indiscriminately across all enhancer types, pinpointing NFE2L2 as the selective lexifier in these cells. Previously, phase separation has been largely attributed to co-activators^21^, and to few sequence-specific TFs such as OCT4^28,77^. Our findings establish NFE2L2 as the sequence-specific driver of transcriptional condensate formation at SEs, thereby directly coupling DNA sequence recognition with enhancer-promoter communication and transcriptional output.

Phase separation has been implicated in large-scale genome compartmentalization^35^, but in the context of enhancer-promoter communication, condensate formation and loop extrusion have traditionally been regarded as distinct processes. Our results reveal that these processes are mechanistically coupled through binding of a lexifier TF at classical enhancers. NFE2L2-depletion increased CTCF and SMC1 occupancy at SE boundaries, resulting in formation of new sub-TADs and loss of enhancer-promoter contacts. We propose that in the absence of NFE2L2-driven transcriptional condensates, loop extrusion stalls, leading to excessive cohesin clogging on chromatin. Thus, in addition to previously described mechanisms controlling cohesin occupancy, such as its ATPase activity^24,78^ and loading and unloading factors^43,45^, we describe a novel mechanism for impaired extrusion dynamics caused by loss of lexifier binding at classical enhancers.

Lexifier TFs play a pivotal role in coupling sequence recognition with condensate formation and productive loop extrusion, maintaining the structural integrity of SEs and the regulatory hierarchy of multipartite enhancer clusters. Despite the predicted formation of several new sub-TADs at the *MYC* locus, MYC expression remained unaffected, suggesting that active *cis*-regulatory elements can mediate clustered interactions that bypass TAD boundaries. Our findings align with recent reports proposing that phase separation acts cooperatively with loop extrusion to fine-tune genome organization^77,79,80^ and that enhancer RNA production at regulatory elements contributes to condensate formation^81^. The “three-way kissing” model^34^ further supports a cooperative role between SEs, CTCF, and condensates in regulating gene expression. Consistent with this interplay, while cohesin and CTCF are essential for TAD formation^40,82,83^, some enhancer-promoter contacts persist after cohesin loss^82,84,85^, suggesting that other mechanisms — such as TF-driven condensates — may sustain regulatory interactions^34^.

Sequence-specific TFs have a well-established role in decoding the regulatory blueprint of the genome by binding promoters and enhancers in a context-specific manner^50,86^, and they can even repurpose silenced repetitive elements to drive aberrant transcription in cancer^87–89^. NFE2L2 typically coordinates oxidative stress responses but is frequently hyperactivated in cancer^90^. TCF7L2, which we found enriched at classical enhancers in GP5d cells, functions as a key effector of Wnt/β-catenin signaling during development and is linked to enhancer activity in colorectal cancer^91^. Here, we show that these TFs have multifaceted roles beyond transcriptional control, contributing to genome integrity and 3D chromatin organization. Our results further suggest that other sequence-specific TFs with similar biochemical properties may act as lexifiers, organizing SE architecture and regulatory hierarchy in a cell type-selective manner. This is supported by the reliance of liver and colon cancer cells on different lexifier TFs and the observation that not all SEs in HepG2 cells are bound by NFE2L2, despite widespread genome reorganization upon its depletion, suggesting that other TFs can function as lexifiers in a locus-specific manner. Together, these findings define an emerging paradigm in which lexifier TFs couple DNA sequence recognition to higher-order genome organization, establishing a framework for understanding enhancer hierarchy and its rewiring in disease.

In conclusion, our study provides a comprehensive framework linking enhancer sequence, TF binding, phase separation, and 3D genome organization. By functionally distinguishing classical enhancers from facilitators within SEs, we establish the deterministic role of classical enhancers and their decoding by specific lexifier TFs such as NFE2L2 in HepG2 cells. These findings highlight the functional heterogeneity within SEs and uncover multifaceted roles for TFs in coordinating transcriptional output and genome architecture. Beyond advancing our understanding of enhancer logic, this work provides a conceptual foundation for exploring how dysregulation of the lexifier–enhancer axis contributes to developmental disorders, cancer, and other diseases driven by aberrant genome regulation.

## Methods

### Data acquisition

All sequencing datasets and corresponding annotation file download links utilized in this study are provided in **Supplementary Table 5**, together with relevant GEO/ENCODE accession numbers.

Human Hg38 blacklisted regions were downloaded from ENCODE^50^ (ENCFF356LFX).

PhastCons100 and PhyloP100 conservation score bigwig files were downloaded from UCSC (https://hgdownload.cse.ucsc.edu/goldenpath).

Hg38 chromosome sizes file was downloaded from UCSC (https://hgdownload.soe.ucsc.edu/goldenpath/).

Motif file for AME analysis was downloaded from JASPAR 2022 (https://jaspar2022.genereg.net/download/data/2022/CORE/JASPAR2022_CORE_verte 607 brates_non-redundant_pfms_meme.txt)

Motif clustering file was downloaded from (https://resources.altius.org/~jvierstra/projects/motif-clustering-v2.0beta/).

A gene annotation GTF file was obtained from Gencode Release v.42, corresponding to the reference chromosomes. This file was converted to BED format using the gtfToBed.sh script. Subsequently, transcription start site (TSS) and gene body BED files were created with a custom script adapted from Lee et al.^92^

### Cell culture

ChIP-seq was performed as previously described^89^ by using the following antibodies for: NFE2L2 (Abcam, ab62352), MED1 (Bethyl Labs, A300-793A), P300 (Santa Cruz Biotechnology, sc-585x), H3K27ac (Diagenode, C15410196), BRD4 (Cell Signaling Technology, 13440S), SMC1 (Bethyl Labs, A300-055A), CTCF (Abcam, ab70303), NIPBL (Bethyl Labs, A301-778A) and WAPL (Proteintech, 16370-1-AP). ChIP-seq was performed by using 2-5 μg of antibody per reaction. HepG2 cells were formaldehyde cross-linked for 10 minutes (min) at room temperature (RT). Sonicated chromatin was centrifuged, and the supernatant was used to immunoprecipitate DNA using Dynal-bead coupled antibodies. Immunoprecipitated DNA was purified and used for ChIP-seq library for Illumina sequencing. The libraries were single-read sequenced on NovaSeq 6000 and NovaSeq X.

### ChIP-seq

ChIP-seq was performed as previously described^89^ by using the following antibodies for: NFE2L2 (Abcam, ab62352), MED1 (Bethyl Labs, A300-793A), P300 (Santa Cruz Biotechnology, sc-585x), H3K27ac (Diagenode, C15410196), BRD4 (Cell Signaling Technology, 13440S), SMC1 (Bethyl Labs, A300-055A), CTCF (Abcam, ab70303), NIPBL (Bethyl Labs, A301-778A) and WAPL (Proteintech, 16370-1-AP). ChIP-seq was performed by using 2-5 μg of antibody per reaction. HepG2 cells were formaldehyde cross-linked for 10 minutes (min) at room temperature (RT). Sonicated chromatin was centrifuged, and the supernatant was used to immunoprecipitate DNA using Dynal-bead coupled antibodies. Immunoprecipitated DNA was purified and used for ChIP-seq library for Illumina sequencing. The libraries were single-read sequenced on NovaSeq 6000 and NovaSeq X.

### ATAC-seq

ATAC-seq library was performed by using 50000 HepG2 cells according to the protocol described earlier^89^. HepG2 cells were washed with ice-cold PBS and were resuspended in 50 µl lysis buffer. The cells were incubated for 10 min on ice. The pellet was resuspended in 2×tagmentation buffer (Illumina kit) and incubated at 37 °C for 30 min. DNA was purified by using MinElute purification kit and eluted in elution buffer. Optimal number of amplification PCR cycles was determined by qPCR. Samples were amplified by using Nextera library preparation kit (Illumina) and sequenced paired end in Novaseq 6000.

### RNA-seq

Total RNA was extracted from HepG2 cells using the RNeasy Mini Kit (Qiagen). RNA-seq libraries were prepared from 500 ng of total RNA using the KAPA Stranded RNA-Seq Kit for Illumina (Roche), following the manufacturer’s protocol. Paired-end sequencing was performed on an Illumina NovaSeq 6000.

### PRO-seq

PRO-seq was performed with minor modifications to previously published protocols^93,94^. HepG2 WT and NFE2L2 KD cells (12 million/sample) were trypsinized, washed with ice-cold PBS, and permeabilized in permeabilization buffer on ice for 5 min. For nuclear run-on, 10 million permeabilized HepG2 cells were mixed with 200,000 Drosophila S2 cells (spike-in) and incubated at 37°C for 5 min in 2× nuclear run-on buffer containing biotin-11-NTPs (Jena Bioscience), SuperaseIN (Thermo Fisher Scientific, AM2694), and 1% sarkosyl (Thero Fisher Scientific, BP234-500).

Total RNA was extracted with TRIzol (Thermo Fisher Scientific, 15596026), hydrolyzed with 1 N NaOH (10 min, ice), and ligated to 3′ adapter (RA3) using T4 RNA ligase (New England Biolabs, M0204S) with 15% PEG-8000 (overnight, 16°C). Streptavidin beads (20 µL/sample; Thermo Fisher Scientific, 15596018) were washed and used to capture biotin-labeled RNA. Beads were washed with high- and low-salt buffers, followed by on-bead 5′ decapping, hydroxyl repair, and 5′ adapter (RA5) ligation (2 h, 16°C).

RNA was extracted, reverse-transcribed with RP1 primer, and PCR cycle number was optimized via test amplification. Final libraries were amplified with TruSeq primers (RP1 and RPI-X), size-selected by agarose gel electrophoresis, and subjected to paired-end sequencing on an Illumina NovaSeq X. The sequences of all primers and RNA adaptors used in PRO-seq library preparation are listed in **Supplementary Table 6**.

### Micro-Capture-C

MCC was performed as previously described^95^. Biotynylated probes for the candidate enhancers and gene promoters were designed by using IDT xGen™ MRD Hyb Panel and ordered from IDT. Six million HepG2-WT and NFE2L2 KD cells per sample, two biological replicates, and two technical replicates for each biological replicate. Cells were crosslinked with 2% formaldehyde, quenched with 125 mM glycine, and digested *in situ* with MNase (10 Kunitz units, NEB M0247S). Chromatin was ligated, and efficiency assessed via Bioanalyzer.

DNA was extracted (Qiagen DNeasy), sonicated to ∼300 bp (Covaris S220), verified by Bioanalyzer, and purified with 1.8× AMPure XP beads. Sonicated libraries were then subjected to end-repair, adaptor ligation and sequencing indices. 200 ng of indexed libraries were pooled (1:1 mass ratio) and hybridized with 5’-biotinylated capture oligos at 55 °C for 24 h. M-270 Streptavidin Dynabeads (Invitrogen 65305) were used to capture the hybridized capture oligonucleotides and processed as per KAPA HyperCap v3.2 (Roche). Libraries were PCR-enriched (18 cycles) and purified with 1.8X AMPure XP beads. Paired-end sequencing was performed on an Illumina NovaSeq X. MCC probes that were used are available in the **Supplementary Table 7**.

### Phase separation experiments

HepG2 (ATCC, HB-8065, male biological origin) and GP5d (Sigma, 95090715, female biological origin) cells were cultured in DMEM (Gibco, 11960085) supplemented with 10% FBS (Gibco, 10270106), 2mM-glutamine (Gibco, 25030024), and 1% penicillin-streptomycin (Gibco,15140122). Drosophila S2 cells (male biological origin) were kind gift from The DNA fragments encoding BRD4-IDR, MED1-IDR, NFE2L2 CDS full length, NFE2L2-IDR, NFE2L2-IDR deleted-CDS were cloned to the pET-45b-mEGFP vector (Addgene #185013) with 6xHis-tagged. The EGFP fusion protein was expressed in *Escherichia coli* BL21(DE3) cells in LB medium. The plasmids from a single clone were extracted and verified by Sanger sequencing. The bacteria were cultured in LB until OD reached 0.6 and incubated with isopropyl-β-D-thiogalactopyranoside (IPTG) for 4 hours. Pellets for GFP fusion protein extraction were collected by using HisPur™ Ni-NTA Purification Kit (Thermo fisher, 88228). The HepG2 nuclear protein was extracted by NE-PER Nuclear Extraction Reagents (Thermo fisher, 78833).

Recombinant IDRs of MED1, BRD4, NFE2L2, and NFE2L2 WT, NFE2L2-IDR deleted-CDS fusion proteins were concentrated and desalted to 15 µM by using Amicon Ultra centrifugal filters (Millipore). Recombinant protein was added to solutions with or without nuclear extract and PEG 8000 (final concentrations, 10%) in buffer (50 mM Tris, pH 7.2, 10% glycerol, and 1 mM DTT). The fusion proteins mixed with nuclear extract were loaded onto a chamber slide and imaged with a fluorescent microscope.

### RT-qPCR

To validate the expression of MCC target genes by RT-qPCR, total RNA was extracted from homozygous deletion clones using the RNeasy Mini Kit (Qiagen). cDNA was synthesized using the PrimeScript™ RT Master Mix (Takara, RR036A). RT-qPCR was then performed in triplicate using the SYBR Green I Master Mix (Roche, 04707516001). Gene expression levels were normalized to GAPDH. Primer sequences for each gene are provided in **Supplementary Table 6.**

### CRISPR enhancer deletion

CRISPR-mediated knockout of classical enhancers and facilitators was carried out as previously described^89^. Two sgRNAs targeting each flank of the enhancer were designed using CRISPOR v.5.2 and synthesized as crRNAs by Integrated DNA Technologies (IDT) (**Supplementary Table 6**). Briefly, equimolar amounts of enhancer-specific crRNAs and ATTO550-labeled tracrRNA (IDT, 1075928) were annealed. The resulting duplexes were complexed with Alt-R CRISPR-Cas9 nuclease (IDT, 1081060; 1000 ng per 200,000 cells) and target-specific sgRNAs (250 ng per 200,000 cells) to form ribonucleoprotein (RNP) complexes. These RNPs were transfected into HepG2 cells using CRISPRMAX (Life Technologies, CMAX000003) following the manufacturer’s instructions. On the following day, ATTO550-positive cells were isolated via FACS (**Extended Data Fig. 10**), and single-cell colonies were expanded to establish clonal enhancer knockout lines. After 2–3 weeks of culture, individual clones were screened for homozygous deletions using rapid DNA lysis (Lucigen, QE0905T) and PCR with primers flanking the expected deletion site (**Supplementary Table 6**). Expression of target genes were analyzed by using RT-qPCR. Primer sequences for each gene are provided in **Supplementary Table 6.**

### CRISPR inhibition of classical enhancers

HepG2 cells were infected with lentivirus expressing dCas9-Mecp2 (Addgene #122205) at MOI = 1. After 48 hours, transduced cells were selected with 2 μg/ml blasticidin for 7 days. sgRNAs targeting the summit of the target enhancers were designed using CRISPOR v.5.2. Forward and reverse oligonucleotides of sgRNAs were synthesized and cloned into the lentiviral vector lentiGuide-Puro (Addgene #52963). Lentivirus packaging was performed, and the resulting sgRNA-expressing lentiviruses (or control lentiviruses expressing non-targeting sgRNA) were used to infect the established dCas9-Mecp2-expressing HepG2 cells at MOI = 1. After 48 hours, transduced cells were selected with 2 μg/ml puromycin for 7 days. Following puromycin selection, total RNA was extracted using the RNeasy Mini Kit (Qiagen). Expression of target genes were compared by using RT-qPCR. sgRNA and primer sequences for each gene are provided in **Supplementary Table 6.**

### CRISPR deletion of NFE2L2

To generate HepG2 NFE2L2 KD cells, two sgRNAs targeting the NFE2L2 exon 2 were designed and individually cloned into the pSpCas9(BB)-2A-Puro (PX459) V2.0 vector (Addgene #62988). HepG2 cells at approximately 70% confluency were co-transfected with both sgRNA-containing PX459 plasmids using Lipofectamine 3000 (Thermo Fisher Scientific). After overnight incubation, the culture medium was replaced with puromycin selection medium (0.8 µg/mL; Sigma-Aldrich). Forty-eight hours post-selection, cells were trypsinized and seeded as single cells into 96-well plates. Clonal expansion was monitored over 9–16 days to ensure monoclonality. Individual clones were subsequently expanded and screened for CRISPR deletion using gel electrophoresis after genomic PCR (**Extended Data Fig. 3b**). The clonal cells lines were screened for NFE2L2 deletion by Western blotting (**Extended Data Fig. 3c**).

### Western blot

Western blots were performed as described earlier^96^. Cells were lysed in RIPA buffer with 1 mM DTT (Thermo Scientific, 20290) and protease inhibitors (Roche, 11873580001). Protein lysates (50 μg per sample) were denatured in 6× SDS-Laemmli buffer at 95 °C for 5 min, separated by SDS-PAGE, and transferred to PVDF membranes. Membranes were blocked in 5% milk/TBST and incubated with primary antibodies: NFE2L2 (Abcam, ab62352, 1:1000) and GAPDH (Santa Cruz Biotechnology, SC-47724, 1:3000). Secondary antibodies (Bio-Rad, 5178-2504 and 5196-2504, 1:5000) were applied, and blots were imaged using Image Studio Lite on the Odyssey CLx imager.

### ChIP-seq analysis

ChIP-seq was analyzed as previously described^89^. In short, sequencing reads were aligned to the hg38/GRCh38 genome using bowtie2 v.2.2.5 (bowtie2 --very-sensitive) (https://bowtie-bio.sourceforge.net/bowtie2/index.shtml). Duplicate reads were identified using Picard v.2.26.3 (http://broadinstitute.github.io/picard/). Duplicate and low-quality reads (Phred < 10) were removed using samtools v.2.30.0 (https://www.htslib.org/). Peak calling was performed using MACS2 v.2.1.1.20160309 (macs2 callPeak --nomodel) (https://pypi.org/project/MACS2/). Reproducible ChIP-seq peaks shared between two replicates identified using bedtools v.2.30.0 (https://github.com/arq5x/bedtools2) and used for downstream analysis. Normalized signal files were created using deepTools v.3.1.3 (bamCoverage --binSize 10 --normalizeUsing RPKM) (https://github.com/deeptools/deepTools). BedGraph files were generated using UCSC bedGraphToBigWig v.377 (https://www.encodeproject.org/software/bedgraphtobigwig/). ChIP-seq signal quantification was performed using bwtool v.1.0 (bwtools summary - header) (https://github.com/CRG-Barcelona/bwtool). Correlation plot for the ChIP-seq replicates is shown in **Extended Data Fig. 8a**.

### ATAC-seq analysis

ATAC-seq was analyzed similar to ChIP-seq by peak calling using MACS2 v.2.2.7.1 (macs2 --nomodel --keep-dup all -g hs). Blacklisted regions were excluded, and coverage tracks were generated and RPKM normalized as described for ChIP-seq data. Correlation plot for the ATAC-seq replicates is shown in **Extended Data Fig. 8a**.

### Genomic STARR-seq analysis

STARR-seq was analyzed as previously described^89^. Reads were aligned to the hg38/GRCh38 genome using Bowtie2 v.2.4.1 (bowtie2 --maxins 1000). Duplicates were marked with Picard v.2.23.4. Samtools v.1.7 was used to filter non-concordant and low-quality reads (-F 1024 -q 20). Peaks were called with MACS2 v.2.2.7.1 (--f BAMPE --g hs) using STARR input as control. RPKM normalized coverage track were generated by using deepTools with the following options (bamCoverage --binSize 10 --normalizeUsing RPKM --extendReads).

### GRO-seq analysis

HepG2 GRO-seq data were obtained from GEO (GSM2428726, SRR5109940). Reads were trimmed using Trim Galore v.0.6.7 (https://www.bioinformatics.babraham.ac.uk/projects/trim_galore/) to remove A-stretches from library preparation. Reads shorter than 25 bp or with quality scores <10 were discarded. Alignment to the hg38 genome was performed using Bowtie2, and strand-specific bigWig files were generated using Samtools v.1.9.

### RNA-seq analysis

RNA-seq reads were aligned to the hg38 reference genome using STAR v.2.7.4a (https://github.com/alexdobin/STAR). Reads were then sorted using samtools and the read counts were quantified using htseq-count v.0.11.2 (https://pypi.org/project/HTSeq/). Low count genes (sum of samples and replicates <10) were filtered out and differential expression analysis was performed using DEseq2 v.1.32.0 (https://www.bioconductor.org/packages//2.13/bioc/html/DESeq2.html). Genes with adjusted p-value < 0.05 and absolute log2FoldChange > 1.5 were considered differentially expressed.

### PRO-seq analysis

PRO-seq analysis was performed by using a custom lab pipeline. In short, paired-end reads were deduplicated using BBtools clumpify (dedupe=t subs=0 unpair=f) (https://sourceforge.net/projects/bbmap/). Adapter and UMI sequences were trimmed with Trim Galore v.0.6.10 (--length 14 --paired --adapter TGGAATTCTCGGGTGCCAAGGAACTCCAGTCAC --gzip --clip_R2 6). Reads were aligned to a combined human (hg38) and Drosophila (dm3) genome using BWA-MEM v.0.7.18, and aligned reads with mapping quality >20 were retained using Samtools.

Aligned BAM files were converted to BED format (bedtools bamToBed) and split into hg38 and dm3 using bedtools intersect. Reads mapping to hg38 genome were used to make strand-specific bedGraph file using bedtools genomeCoverageBed, followed by RPM normalization. Final signal tracks were converted to bigWig using UCSC bedGraphToBigWig.

PRO-seq signal differences between classical enhancers and facilitators in WT and KD conditions were quantified using bwtool (bwtool summary -header -fill=0). The signal was quantified for a region centered on the MED1 enhancer peak, extended 250 base pairs in both directions.

Fox PI analysis, promoter-proximal counts (PPC; defined as 50 bp upstream to 150 bp downstream of the TSS) and gene body counts (GBC; defined as 200 bp downstream of the TSS to 100 bp upstream of the TES) were extracted from RPM-normalized bedGraph files using bedtools coverage v.2.30.0. PI was calculated as the ratio of PPC to GBC. For genes with multiple isoforms, the transcript with the highest sequence length-normalized GBC was selected. PI values for upregulated and downregulated genes were compared between WT and NFE2L2 KD cells using a two-sided paired t-test.

### ChIA-PET analysis

Processed ChIA-PET data for POLR2A was obtained from ENCODE (ENCFF364UNM). The number of interactions at classical enhancers and facilitators quantified using countOverlaps function from GenomicRanges (https://bioconductor.org/packages/release/bioc/html/GenomicRanges.html) R package. The overlapping counts for either up- or downstream anchors of the loops and added a pseudo count of one before log2 transformation. P-value was calculated using unpaired Wilcoxon test.

### Micro-Capture-C analysis

MCC analysis was performed as described in Hamley et al.^95^. Sequencing adapters were trimmed using Trim Galore v.0.6.7, and overlapping reads were merged with FLASH v.2.2.00 (https://www.cbcb.umd.edu/software/flash). Reads were mapped to guide sequences using BLAT v.35 (-minScore=20 -minIdentity=5 -maxIntron=10000 - tileSize=11) (http://www.soe.ucsc.edu/~kent), then separated by probe using the MCC_spliter.pl script. Probe-specific reads were aligned to the hg38 genome using Bowtie2 (bowtie2 -X 1000). Ligation junctions were defined using MCC_analyser.pl script.

Replicates were pooled, and scaling factors were calculated to normalize KD samples to WT depth. Read counts per viewpoint were obtained using Samtools (view -c), and scaling ratios (WT/KD) were applied using deepTools bamCoverage (--scaleFactor <factor> --binSize 1) to generate depth-normalized bigWig files for plotting. To identify top WT contacts per viewpoint, peaks were called with MACS2 (--keep-dup all --nolambda --extsize 100 --nomodel). Peaks were ranked by –log₁₀(q-value), and the top 10 non-viewpoint summits on the same chromosome were selected as the strongest WT interactions.

To quantify ligation junction interactions, we first extracted reads located within ±500 kb of each viewpoint and identified interaction peaks. Each peak was then extended by ±500 bp from its summit, and the number of ligation junction reads within these extended regions was counted. Enhancer sites were annotated using H3K27ac ChIP-seq datasets, respectively. Peaks overlapping their own corresponding viewpoints were excluded.

### Super-enhancer calling

SEs were called using Rank Ordering of Super-enhancers (ROSE) algorithm v.0.1 (https://bitbucket.org/young_computation/rose/src/master/)^9^ and H3K27ac ChIP-seq. Candidate enhancer sites were defined as reproducible peaks from two replicates, and H3K27ac signal was taken from the first replicate using input as a control with options (-s 12500 -t 2500).

### Defining classical and facilitator enhancers

To classify enhancers within SEs, we used STARR-seq and MED1 ChIP-seq peaks. Due to the broad distribution of H3K27ac signals, MED1 peaks were used to define the classical enhancers and facilitators within SEs. First, all enhancers were categorized as either within SEs or outside SEs based on overlap with SE regions using bedtools v.2.30.0. Enhancers within SEs were further classified as classical enhancers if they overlapped STARR-seq peaks, and as facilitators if they did not. The summit of each classical enhancer was defined by the STARR-seq peak summit, while the summit of each facilitator was defined by the corresponding MED1 ChIP-seq peak summit.

### Motif analysis and affinity prediction

Motif analysis was performed as described in Karttunen et al.^89^ using AME v.5.0.2 from the MEME Suite^53^ (ame --control --shuffle). TF binding affinities were predicted using the tRap R package (https://github.com/matthuska/tRap) with human core motifs PWMs from the JASPAR^97^. The NFE2L2 PWM (MA0150.1) was manually added. Regions shorter than the maximum motif length (<33 bp) were excluded. Mean affinity scores were calculated for each motif across core enhancers (classical enhancers) and control distal enhancers (facilitators). Then motifs were assigned into clusters according to Viestra et al.^54^ and mean affinities were computed per cluster for both classical enhancers and facilitators. A representative motif was from each cluster was used to label the cluster.

### Lenti-MPRA analysis

Processed lenti-MPRA datasets were downloaded from Agarwal et al.^51^. Each lenti-MPRA probe was extended by ±200 base pairs. Overlaps between the extended lenti-MPRA probe and enhancer sites were identified using bedtools intersect. Violin plots showing the distribution of the log₂(RNA/DNA) ratios for lenti-MPRA probes were generated in R using the ggplot2 (https://cran.r-project.org/web/packages/ggplot2/index.html) and ggpubr (https://cran.r-project.org/web/packages/ggpubr/index.html) packages. Statistical comparisons between groups were performed using the unpaired Wilcoxon rank-sum test.

### Polymer simulations and analysis

We used MoDLE^59^ to simulate 3D genome organisation for HepG2 WT and NFE2L2 KD cells by using ChIP-seq data. First, CTCF binding sites were predicted genome-wide using MAST v.5.5.8 from the MEME Suite^98^ using the CTCF motif. Predicted CTCF-binding sites were then filtered to retain only those overlapping with CTCF ChIP-seq peaks. Filtered sites were used by MoDLE to generate a BED file of extrusion barriers, incorporating ChIP-seq peak signal as average occupancy. MoDLE then simulated loop extrusion dynamics to generate a contact matrix, output as a .cool file representing predicted chromatin interactions.

Simulated contact maps were analyzed using HiCExplorer v.3.7.3^99^. Matrices at 20 kb resolution were bias-corrected with hicCorrectMatrix (--filterThreshold -1.5 5 -- correctionMethod ICE) and normalized to equal depth using hicNormalize (-normalize smallest). TADs and insulation scores were identified using hicFindTADs with FDR correction.

### Data visualization and statistical analysis

Statistical analyses were done using R v.4.3 and GraphPad prism v.10. Boxplots and violin plots were plotted with ggplot2. Barplots were plotted with GraphPad prism. Motif heatmap was plotted with ComplexHeatmap v.2.22.0. Average profile plots and heatmaps were plotted using deepTools.

Genomic snapshots were plotted with spark.py v.2.6.2 script using option -sm 10 (**Fig. 3**) or -sm 25 (**Fig. 1 and 5, and Extended Data Fig. 1, 2 and 4**). *In silico* snapshots were plotted with HiGlass v.1.1.0 in 5 kb resolution (https://github.com/higlass/higlass) for the 3D contacts and with IGV for the signal tracks (https://igv.org/) and spark.py for gene and SE tracks. MCC snapshots were plotted using pyGenomeTracks^100^ v.3.9. Heatmaps were generated using RPKM-normalized bigWig files from pooled replicates. Heatmaps were created using computeMatrix (scale-regions --skipZeros) and PlotHeatmap from deepTools and the SE regions were scaled to a uniform size of 20 kb. For PRO-seq, the average profile plot was created using (computeMatrix reference-point --missingDataAsZero) and plotProfile.

### Statistics and reproducibility

RNA-seq were performed with three biological replicates. ChIP-seq and ATAC-seq was performed with two biological replicates. The statistical tests used are described in the methods section and respective figure legends.

## Data availability

Sequencing data have been deposited at ENA as PRJEB100961 and are publicly available as of the date of publication.

UCSC genome browser tracks are available at: https://genome.ucsc.edu/s/villet/Tiusanen_Patel_et_al_2025.

Previously published sequencing datasets and annotation links utilized in this study are provided in **Supplementary Table 5**. Previously published data are available under the following accession code: GEO: GSE180158, GSM2428726, GSE180158, GSE254242. ENCODE: ENCFF920MZO, ENCFF655BEL, ENCFF000PIE, ENCFF000PHU, ENCFF000XTR, ENCFF000XTQ, ENCFF000XUL, ENCFF000XUK, ENCFF128UUS, ENCFF015SPJ, ENCFF492CBJ, ENCFF807WOU, ENCFF416JVM, ENCFF382VQI, ENCFF364UNM. Processed lenti-MPRA probe data for HepG2 cells was downloaded from ENCODE: ENCFF774DYO.

## Code availability

This paper does not report original code.

## Author Contributions

BS conceptualized and supervised the study. Experiments were performed by DP, JX, CX, LF, SD, and data analysis was done by VT, DP, SC, JY. MM and SP helped with MCC experiments and analysis. PP and EP helped with scientific discussion and presentation of the data. DP, VT and BS wrote the manuscript with contributions from all authors.

## Acknowledgement

We thank Norwegian Sequencing Centre, Oslo, Norway and HiLIFE research infrastructure for FIMM NGS Genomics laboratory at the University of Helsinki for sequencing services. We thank the Center for Scientific Computing (CSC), Finland, for the computational infrastructure. B.S. was supported by Research Council of Norway (187615), Helse Sør-Øst, University of Oslo through the Norwegian Centre for Molecular Biosciences and Medicine (NCMBM) (to Sahu group), Norwegian Cancer Society (274630), South Eastern Norway Health Authority (HSØ) (2025083), Research Council of Finland (320114), Sigrid Jusélius Foundation, Jane and Aatos Erkko Foundation. BS and EP were supported by iCAN Digital Precision Cancer Medicine Flagship (320185). Research Council of Finland supported P.P. (356021) and J.X. (342594). S.D. was supported by the iCANPOD postdoctoral program through the iCANDOC doctoral pilot in precision cancer medicine. V.T. was supported by doctoral program of Integrative Life Sciences, University of Helsinki. We thank Veera Erkkilä, Pinja Perkkiö and other members of Sahu group for technical assistance and scientific discussion.

## References

1 Dekker, J. & Mirny, L. A. The chromosome folding problem and how cells solve it. Cell 187, 6424–6450 (2024). 10.1016/j.cell.2024.10.026

2 Grubert, F. et al. Landscape of cohesin-mediated chromatin loops in the human genome. Nature 583, 737–743 (2020). 10.1038/s41586-020-2151-x

3 Long, H. K., Prescott, S. L. & Wysocka, J. Ever-Changing Landscapes: Transcriptional Enhancers in Development and Evolution. Cell 167, 1170–1187 (2016). 10.1016/j.cell.2016.09.018

4 Spitz, F. & Furlong, E. E. Transcription factors: from enhancer binding to developmental control. Nat Rev Genet 13, 613–626 (2012). 10.1038/nrg3207

5 Lambert, S. A. et al. The Human Transcription Factors. Cell 172, 650–665 (2018). 10.1016/j.cell.2018.01.029

6 Blayney, J. W. et al. Super-enhancers include classical enhancers and facilitators to fully activate gene expression. Cell 186, 5826–5839 e5818 (2023). 10.1016/j.cell.2023.11.030

7 Blobel, G. A., Higgs, D. R., Mitchell, J. A., Notani, D. & Young, R. A. Testing the super-enhancer concept. Nat Rev Genet 22, 749–755 (2021). 10.1038/s41576-021-00398-w

8 Hnisz, D. et al. Super-enhancers in the control of cell identity and disease. Cell 155, 934–947 (2013). 10.1016/j.cell.2013.09.053

9 Whyte, W. A. et al. Master transcription factors and mediator establish super-enhancers at key cell identity genes. Cell 153, 307–319 (2013). 10.1016/j.cell.2013.03.035

10 Arnold, C. D. et al. Genome-wide quantitative enhancer activity maps identified by STARR-seq. Science 339, 1074–1077 (2013). 10.1126/science.1232542

11 Sahu, B. et al. Sequence determinants of human gene regulatory elements. Nat Genet 54, 283–294 (2022). 10.1038/s41588-021-01009-4

12 Hay, D. et al. Genetic dissection of the alpha-globin super-enhancer in vivo. Nat Genet 48, 895–903 (2016). 10.1038/ng.3605

13 Huang, J. et al. Dissecting super-enhancer hierarchy based on chromatin interactions. Nat Commun 9, 943 (2018). 10.1038/s41467-018-03279-9

14 Shin, H. Y. et al. Hierarchy within the mammary STAT5-driven Wap super-enhancer. Nat Genet 48, 904–911 (2016). 10.1038/ng.3606

15 Thomas, H. F. et al. Temporal dissection of an enhancer cluster reveals distinct temporal and functional contributions of individual elements. Mol Cell 81, 969–982 e913 (2021). 10.1016/j.molcel.2020.12.047

16 Chong, S. et al. Imaging dynamic and selective low-complexity domain interactions that control gene transcription. Science 361 (2018). 10.1126/science.aar2555

17 Kato, M. et al. Cell-free formation of RNA granules: low complexity sequence domains form dynamic fibers within hydrogels. Cell 149, 753–767 (2012). 10.1016/j.cell.2012.04.017

18 Larson, A. G. et al. Liquid droplet formation by HP1alpha suggests a role for phase separation in heterochromatin. Nature 547, 236–240 (2017). 10.1038/nature22822

19 Lin, Y., Protter, D. S., Rosen, M. K. & Parker, R. Formation and Maturation of Phase-Separated Liquid Droplets by RNA-Binding Proteins. Mol Cell 60, 208–219 (2015). 10.1016/j.molcel.2015.08.018

20 Pei, G., Lyons, H., Li, P. & Sabari, B. R. Transcription regulation by biomolecular condensates. Nat Rev Mol Cell Biol 26, 213–236 (2025). 10.1038/s41580-024-00789-x

21 Sabari, B. R. et al. Coactivator condensation at super-enhancers links phase separation and gene control. Science 361 (2018). 10.1126/science.aar3958

22 Sabari, B. R., Hyman, A. A. & Hnisz, D. Functional specificity in biomolecular condensates revealed by genetic complementation. Nat Rev Genet 26, 279–290 (2025). 10.1038/s41576-024-00780-4

23 Davidson, I. F. et al. DNA loop extrusion by human cohesin. Science 366, 1338–1345 (2019). 10.1126/science.aaz3418

24 Kim, Y., Shi, Z., Zhang, H., Finkelstein, I. J. & Yu, H. Human cohesin compacts DNA by loop extrusion. Science 366, 1345–1349 (2019). 10.1126/science.aaz4475

25 Mirny, L. & Dekker, J. Mechanisms of Chromosome Folding and Nuclear Organization: Their Interplay and Open Questions. Cold Spring Harb Perspect Biol 14 (2022). 10.1101/cshperspect.a040147

26 Dignon, G. L., Best, R. B. & Mittal, J. Biomolecular Phase Separation: From Molecular Driving Forces to Macroscopic Properties. Annu Rev Phys Chem 71, 53–75 (2020). 10.1146/annurev-physchem-071819-113553

27 Boehning, M. et al. RNA polymerase II clustering through carboxy-terminal domain phase separation. Nat Struct Mol Biol 25, 833–840 (2018). 10.1038/s41594-018-0112-y

28 Boija, A. et al. Transcription Factors Activate Genes through the Phase-Separation Capacity of Their Activation Domains. Cell 175, 1842–1855 e1816 (2018). 10.1016/j.cell.2018.10.042

29 Cho, W. K. et al. Mediator and RNA polymerase II clusters associate in transcription-dependent condensates. Science 361, 412–415 (2018). 10.1126/science.aar4199

30 Mensah, M. A. et al. Aberrant phase separation and nucleolar dysfunction in rare genetic diseases. Nature 614, 564–571 (2023). 10.1038/s41586-022-05682-1

31 Ahn, J. H. et al. The phenylalanine-and-glycine repeats of NUP98 oncofusions form condensates that selectively partition transcriptional coactivators. Mol Cell 85, 708–725 e709 (2025). 10.1016/j.molcel.2024.12.026

32 Lyons, H. et al. RNA polymerase II partitioning is a shared feature of diverse oncofusion condensates. Cell (2025). 10.1016/j.cell.2025.04.002

33 Hnisz, D., Shrinivas, K., Young, R. A., Chakraborty, A. K. & Sharp, P. A. A Phase Separation Model for Transcriptional Control. Cell 169, 13–23 (2017). 10.1016/j.cell.2017.02.007

34 Du, M. et al. Direct observation of a condensate effect on super-enhancer controlled gene bursting. Cell 187, 331–344 e317 (2024). 10.1016/j.cell.2023.12.005

35 Ganji, M. et al. Real-time imaging of DNA loop extrusion by condensin. Science 360, 102–105 (2018). 10.1126/science.aar7831

36 Sanborn, A. L. et al. Chromatin extrusion explains key features of loop and domain formation in wild-type and engineered genomes. Proc Natl Acad Sci U S A 112, E6456–6465 (2015). 10.1073/pnas.1518552112

37 Fudenberg, G. et al. Formation of Chromosomal Domains by Loop Extrusion. Cell Rep 15, 2038–2049 (2016). 10.1016/j.celrep.2016.04.085

38 Rao, S. S. et al. A 3D map of the human genome at kilobase resolution reveals principles of chromatin looping. Cell 159, 1665–1680 (2014). 10.1016/j.cell.2014.11.021

39 Dixon, J. R. et al. Topological domains in mammalian genomes identified by analysis of chromatin interactions. Nature 485, 376–380 (2012). 10.1038/nature11082

40 Nora, E. P. et al. Targeted Degradation of CTCF Decouples Local Insulation of Chromosome Domains from Genomic Compartmentalization. Cell 169, 930–944 e922 (2017). 10.1016/j.cell.2017.05.004

41 Rowley, M. J. & Corces, V. G. Organizational principles of 3D genome architecture. Nat Rev Genet 19, 789–800 (2018). 10.1038/s41576-018-0060-8

42 Conte, M. et al. Loop-extrusion and polymer phase-separation can co-exist at the single-molecule level to shape chromatin folding. Nat Commun 13, 4070 (2022). 10.1038/s41467-022-31856-6

43 Ciosk, R. et al. Cohesin’s binding to chromosomes depends on a separate complex consisting of Scc2 and Scc4 proteins. Mol Cell 5, 243–254 (2000). 10.1016/s1097-2765(00)80420-7

44 Hansen, A. S. CTCF as a boundary factor for cohesin-mediated loop extrusion: evidence for a multi-step mechanism. Nucleus 11, 132–148 (2020). 10.1080/19491034.2020.1782024

45 Kueng, S. et al. Wapl controls the dynamic association of cohesin with chromatin. Cell 127, 955–967 (2006). 10.1016/j.cell.2006.09.040

46 Liu, N. Q. et al. WAPL maintains a cohesin loading cycle to preserve cell-type-specific distal gene regulation. Nat Genet 53, 100–109 (2021). 10.1038/s41588-020-00744-4

47 Karpinska, M. A. et al. CTCF depletion decouples enhancer-mediated gene activation from chromatin hub formation. Nat Struct Mol Biol 32, 1268–1281 (2025). 10.1038/s41594-025-01555-z

48 Karpinska, M. A. & Oudelaar, A. M. The role of loop extrusion in enhancer-mediated gene activation. Curr Opin Genet Dev 79, 102022 (2023). 10.1016/j.gde.2023.102022

49 Bouvy-Liivrand, M. et al. Analysis of primary microRNA loci from nascent transcriptomes reveals regulatory domains governed by chromatin architecture. Nucleic Acids Res 45, 9837–9849 (2017). 10.1093/nar/gkx680

50 Consortium, E. P. An integrated encyclopedia of DNA elements in the human genome. Nature 489, 57–74 (2012). 10.1038/nature11247

51 Agarwal, V. et al. Massively parallel characterization of transcriptional regulatory elements. Nature 639, 411–420 (2025). 10.1038/s41586-024-08430-9

52 Roider, H. G., Kanhere, A., Manke, T. & Vingron, M. Predicting transcription factor affinities to DNA from a biophysical model. Bioinformatics 23, 134–141 (2007). 10.1093/bioinformatics/btl565

53 McLeay, R. C. & Bailey, T. L. Motif Enrichment Analysis: a unified framework and an evaluation on ChIP data. BMC Bioinformatics 11, 165 (2010). 10.1186/1471-2105-11-165

54 Vierstra, J. et al. Global reference mapping of human transcription factor footprints. Nature 583, 729–736 (2020). 10.1038/s41586-020-2528-x

55 Abramson, J. et al. Accurate structure prediction of biomolecular interactions with AlphaFold 3. Nature 630, 493–500 (2024). 10.1038/s41586-024-07487-w

56 Tunyasuvunakool, K. et al. Highly accurate protein structure prediction for the human proteome. Nature 596, 590–596 (2021). 10.1038/s41586-021-03828-1

57 Rostam, N. et al. CD-CODE: crowdsourcing condensate database and encyclopedia. Nat Methods 20, 673–676 (2023). 10.1038/s41592-023-01831-0

58 Meszaros, B., Erdos, G. & Dosztanyi, Z. IUPred2A: context-dependent prediction of protein disorder as a function of redox state and protein binding. Nucleic Acids Res 46, W329–W337 (2018). 10.1093/nar/gky384

59 Rossini, R., Kumar, V., Mathelier, A., Rognes, T. & Paulsen, J. MoDLE: high-performance stochastic modeling of DNA loop extrusion interactions. Genome Biol 23, 247 (2022). 10.1186/s13059-022-02815-7

60 Dave, K. et al. Mice deficient of Myc super-enhancer region reveal differential control mechanism between normal and pathological growth. Elife 6 (2017). 10.7554/eLife.23382

61 Zimmerman, M. W. et al. MYC Drives a Subset of High-Risk Pediatric Neuroblastomas and Is Activated through Mechanisms Including Enhancer Hijacking and Focal Enhancer Amplification. Cancer Discov 8, 320–335 (2018). 10.1158/2159-8290.CD-17-0993

62 Hua, P. et al. Defining genome architecture at base-pair resolution. Nature 595, 125–129 (2021). 10.1038/s41586-021-03639-4

63 Arnold, C. D. et al. Quantitative genome-wide enhancer activity maps for five Drosophila species show functional enhancer conservation and turnover during cis-regulatory evolution. Nat Genet 46, 685–692 (2014). 10.1038/ng.3009

64 Arnold, C. D. et al. Genome-wide assessment of sequence-intrinsic enhancer responsiveness at single-base-pair resolution. Nat Biotechnol 35, 136–144 (2017). 10.1038/nbt.3739

65 Liu, Y. et al. Functional assessment of human enhancer activities using whole-genome STARR-sequencing. Genome Biol 18, 219 (2017). 10.1186/s13059-017-1345-5

66 Deng, C. et al. Massively parallel characterization of regulatory elements in the developing human cortex. Science 384, eadh0559 (2024). 10.1126/science.adh0559

67 de Almeida, B. P., Reiter, F., Pagani, M. & Stark, A. DeepSTARR predicts enhancer activity from DNA sequence and enables the de novo design of synthetic enhancers. Nat Genet 54, 613–624 (2022). 10.1038/s41588-022-01048-5

68 Barbadilla-Martinez, L., Klaassen, N., van Steensel, B. & de Ridder, J. Predicting gene expression from DNA sequence using deep learning models. Nat Rev Genet 26, 666–680 (2025). 10.1038/s41576-025-00841-2

69 Avsec, Z. et al. Effective gene expression prediction from sequence by integrating long-range interactions. Nat Methods 18, 1196–1203 (2021). 10.1038/s41592-021-01252-x

70 Zhou, J. et al. Deep learning sequence-based ab initio prediction of variant effects on expression and disease risk. Nat Genet 50, 1171–1179 (2018). 10.1038/s41588-018-0160-6

71 de Almeida, B. P. et al. Targeted design of synthetic enhancers for selected tissues in the Drosophila embryo. Nature 626, 207–211 (2024). 10.1038/s41586-023-06905-9

72 Taskiran, II et al. Cell-type-directed design of synthetic enhancers. Nature 626, 212–220 (2024). 10.1038/s41586-023-06936-2

73 Li, J. et al. Modeling and designing enhancers by introducing and harnessing transcription factor binding units. Nat Commun 16, 1469 (2025). 10.1038/s41467-025-56749-2

74 Ruder, W. C., Lu, T. & Collins, J. J. Synthetic biology moving into the clinic. Science 333, 1248–1252 (2011). 10.1126/science.1206843

75 Brosh, R. et al. Synthetic regulatory genomics uncovers enhancer context dependence at the Sox2 locus. Mol Cell 83, 1140–1152 e1147 (2023). 10.1016/j.molcel.2023.02.027

76 Jindal, G. A. & Farley, E. K. Enhancer grammar in development, evolution, and disease: dependencies and interplay. Dev Cell 56, 575–587 (2021). 10.1016/j.devcel.2021.02.016

77 Wang, J. et al. Phase separation of OCT4 controls TAD reorganization to promote cell fate transitions. Cell Stem Cell 28, 1868–1883 e1811 (2021). 10.1016/j.stem.2021.04.023

78 Elbatsh, A. M. O. et al. Cohesin Releases DNA through Asymmetric ATPase-Driven Ring Opening. Mol Cell 61, 575–588 (2016). 10.1016/j.molcel.2016.01.025

79 Lee, R. et al. CTCF-mediated chromatin looping provides a topological framework for the formation of phase-separated transcriptional condensates. Nucleic Acids Res 50, 207–226 (2022). 10.1093/nar/gkab1242

80 Ryu, J. K. et al. Bridging-induced phase separation induced by cohesin SMC protein complexes. Sci Adv 7 (2021). 10.1126/sciadv.abe5905

81 Henninger, J. E. et al. RNA-Mediated Feedback Control of Transcriptional Condensates. Cell 184, 207–225 e224 (2021). 10.1016/j.cell.2020.11.030

82 Rao, S. S. P. et al. Cohesin Loss Eliminates All Loop Domains. Cell 171, 305–320 e324 (2017). 10.1016/j.cell.2017.09.026

83 Schwarzer, W. et al. Two independent modes of chromatin organization revealed by cohesin removal. Nature 551, 51–56 (2017). 10.1038/nature24281

84 Hsieh, T. S. et al. Enhancer-promoter interactions and transcription are largely maintained upon acute loss of CTCF, cohesin, WAPL or YY1. Nat Genet 54, 1919–1932 (2022). 10.1038/s41588-022-01223-8

85 Jeong, D., Shi, G., Li, X. & Thirumalai, D. Structural basis for the preservation of a subset of topologically associating domains in interphase chromosomes upon cohesin depletion. Elife 12 (2024). 10.7554/eLife.88564

86 Weidemuller, P., Kholmatov, M., Petsalaki, E. & Zaugg, J. B. Transcription factors: Bridge between cell signaling and gene regulation. Proteomics 21, e2000034 (2021). 10.1002/pmic.202000034

87 Chuong, E. B., Elde, N. C. & Feschotte, C. Regulatory activities of transposable elements: from conflicts to benefits. Nat Rev Genet 18, 71–86 (2017). 10.1038/nrg.2016.139

88 Fueyo, R., Judd, J., Feschotte, C. & Wysocka, J. Roles of transposable elements in the regulation of mammalian transcription. Nat Rev Mol Cell Biol 23, 481–497 (2022). 10.1038/s41580-022-00457-y

89 Karttunen, K. et al. Transposable elements as tissue-specific enhancers in cancers of endodermal lineage. Nat Commun 14, 5313 (2023). 10.1038/s41467-023-41081-4

90 DeNicola, G. M. et al. Oncogene-induced Nrf2 transcription promotes ROS detoxification and tumorigenesis. Nature 475, 106–109 (2011). 10.1038/nature10189

91 Del Bosque-Plata, L., Hernandez-Cortes, E. P. & Gragnoli, C. The broad pathogenetic role of TCF7L2 in human diseases beyond type 2 diabetes. J Cell Physiol 237, 301–312 (2022). 10.1002/jcp.30581

92 Lee, I. et al. Simultaneous profiling of chromatin accessibility and methylation on human cell lines with nanopore sequencing. Nat Methods 17, 1191–1199 (2020). 10.1038/s41592-020-01000-7

93 Judd, J. et al. A rapid, sensitive, scalable method for Precision Run-On sequencing (PRO-seq). bioRxiv, 2020.2005.2018.102277 (2020). 10.1101/2020.05.18.102277

94 Vihervaara, A. et al. Transcriptional response to stress is pre-wired by promoter and enhancer architecture. Nat Commun 8, 255 (2017). 10.1038/s41467-017-00151-0

95 Hamley, J. C., Li, H., Denny, N., Downes, D. & Davies, J. O. J. Determining chromatin architecture with Micro Capture-C. Nat Protoc 18, 1687–1711 (2023). 10.1038/s41596-023-00817-8

96 Patel, D., Tiusanen, V., Karttunen, K., Pihlajamaa, P. & Sahu, B. Cancer cell type-specific derepression of transposable elements by inhibition of chromatin modifier enzymes. Commun Biol 8, 992 (2025). 10.1038/s42003-025-08413-0

97 Khan, A. et al. JASPAR 2018: update of the open-access database of transcription factor binding profiles and its web framework. Nucleic Acids Res 46, D260–D266 (2018). 10.1093/nar/gkx1126

98 Bailey, T. L., Johnson, J., Grant, C. E. & Noble, W. S. The MEME Suite. Nucleic Acids Res 43, W39–49 (2015). 10.1093/nar/gkv416

99 Ramirez, F. et al. High-resolution TADs reveal DNA sequences underlying genome organization in flies. Nat Commun 9, 189 (2018). 10.1038/s41467-017-02525-w

100 Lopez-Delisle, L. et al. pyGenomeTracks: reproducible plots for multivariate genomic datasets. Bioinformatics 37, 422–423 (2021). 10.1093/bioinformatics/btaa692

